# Multi-stage physics-informed neural networks for JAK–STAT5 signaling and ultradian insulin–glucose dynamics: latent-species identifiability and suppression of parameter-induced divergence

**DOI:** 10.64898/2026.06.13.728660

**Authors:** Jie Deng, Xinyu Zhang, Xuchang Zhang, Xing Yang

**Affiliations:** School of Basic Medical Sciences, Guilin Medical University, Guilin, Guangxi, China; School of Materials Science and Engineering, Guilin University of Electronic Technology, Guilin, Guangxi, China

**Keywords:** physics-informed neural networks, diffusion–reaction equations, JAK–STAT5 signaling, ultradian insulin–glucose oscillations, latent-species identifiability, parameter-induced trajectory divergence

## Abstract

Coupled diffusion–reaction partial differential equations (PDEs) describe biochemical network dynamics but are difficult to solve for realistic multi-species systems without combining mechanism and data. We present a multi-stage physics-informed neural network (PINN) for multi-species diffusion–reaction PDEs and apply it to two ordinary-differential-equation (ODE) reference systems: the Boehm et al. JAK–STAT5 signaling pathway and the Sturis ultradian insulin–glucose model. For STAT5 we pose a latent-species *identifiability* test: given sparse observations of eight species, a ten-species model that retains two deliberately withheld but mechanistically standard components—an active receptor–JAK complex and the SOCS negative-feedback inhibitor—recovers the reference trajectory and reduces mean root-mean-square error 3.1-fold relative to an eight-species model that omits them, whereas a PDE-only solution without data anchoring diverges. Because the reference is itself ODE-generated, this demonstrates identifiability against synthetic data, not the discovery of new biology. For the insulin–glucose model the same framework reproduces the ∼120-minute oscillation to 1.0% mean relative error as a benchmark on a stiff, multi-timescale oscillator; its spatial dimension is treated as a numerical construct, not a physical transport setting. A Lyapunov analysis of the STAT5 ODE returns a maximal exponent statistically indistinguishable from zero (*λ*_max_ ≈ 3.61 × 10^−5^ min^−1^, 5/8 trials positive; Lyapunov time ∼1.9 × 10^4^ min, far exceeding the 240–720 min horizon), so the system is effectively non-chaotic and the relevant instability is a bounded, parameter-induced trajectory divergence. Anchoring the solution to baseline data suppresses this divergence, with the reduction growing monotonically with sampling density—from ∼15–19% at eight time points to ∼88–97% at sixty-four, depending on perturbation magnitude. The framework thus offers a data-anchored route to latent-species identifiability and divergence suppression in biochemical ODE/PDE systems, demonstrated here against synthetic reference data.

Inside cells, a three-dimensional chemistry of diffusing, reacting molecules drives signaling and rhythm—dynamics that, for realistic networks, strain conventional solvers. Here a multi-stage physics-informed neural network—machine learning constrained by the governing equations—solves stiff, multi-species reaction systems from sparse data. In the JAK–STAT5 signaling pathway, a model that retains two standard but unobserved components (an active receptor complex and a negative-feedback brake) recovers a reference trajectory that a reduced model cannot—a controlled test of whether sparse data can pin down withheld pecies, not a claim of new biology. The same framework reproduces the roughly two-hour insulin–glucose rhythm to within 1% as a benchmark on a stiff oscillator. And anchoring the solution to a few dozen baseline measurements collapses parameter-induced trajectory divergence, turning a parametrically sensitive simulation into a stable one. Where mechanism and data meet, sparse measurements can constrain the structure a model would otherwise leave undetermined.

## 1. Introduction

The dynamics of living systems are encoded in the spatiotemporal evolution of interacting biochemical species—molecules that diffuse, react, and self-organize across scales spanning nanometers to organs and milliseconds to hours [1, 2]. Capturing these dynamics quantitatively is among the most consequential open problems in computational biophysics, and an intrinsically interdisciplinary one: it requires the physics of three-dimensional transport to be reconciled with the incomplete, sparsely sampled knowledge that characterizes real biological systems. Signaling cascades such as the JAK–STAT5 pathway and ultradian rhythms in insulin–glucose regulation exemplify two classes of nonlinear biochemical phenomena whose faithful description demands models that simultaneously resolve spatial transport, multi-species coupling, and intrinsic temporal complexity. Yet a fundamental tension persists in the field: mechanistic models built from first-principles kinetics require parametric and structural knowledge that is rarely complete, while data-driven approaches—however flexible—sacrifice the physical interpretability and extrapolative power that biological inference demands. Physics-informed neural networks (PINNs), which embed partial differential equation (PDE) residuals into neural network optimization [3], sit at the interface between these traditions: they let mechanistic transport equations and observational data jointly constrain a single solution. This positioning is what makes them suited to the sparse-data regimes ubiquitous in experimental biology—where measurements are few, noisy and unevenly sampled—and where a central goal is to constrain, and where possible identify, the unmeasured structure behind the data. Nevertheless, their application to realistic, multi-species biochemical systems governed by three-dimensional diffusion–reaction PDEs remains largely unexplored, and questions regarding the identifiability of unobserved species, the stability of solutions under parametric perturbation, and the adequacy of current modeling paradigms remain open.

The past five years have seen rapid progress in applying physics-informed and physics-encoded learning to reaction–diffusion systems. Lagergren et al. [4] introduced biologically-informed neural networks (BINNs) that simultaneously infer PDE solutions and unknown nonlinear reaction–diffusion terms from sparse cell-migration data, recovering hidden delay mechanisms that eluded conventional modeling. Rao et al. [5] hard-coded reaction–diffusion structure into recurrent convolutional architectures (PeRCNN), demonstrating superior robustness to noise and sparsity relative to standard PINNs, while Chen and Liu [6] coupled PINNs with sparse regression to discover closed-form PDEs—including spiral-wave reaction–diffusion systems—from scarce observations, establishing that physics constraints can regularize learning in chaotic spatiotemporal regimes. Bezekçi [7] further showed that exponential residual schemes stabilize PINNs for stiff convection–reaction–diffusion problems. Karniadakis et al. [8] provided a comprehensive review of the emerging field, highlighting both the promise and the outstanding challenges of physics-informed machine learning. In the metabolic context, Vandvajdi et al. [9] introduced latent variables into a universal PINN framework and, working with experimental glucose–lactate measurements in glioblastoma, used them to uncover signatures of unmodeled metabolic dynamics from real data—a validated demonstration that physics-informed networks can expose unobserved biochemical structure in measurements. The present study is complementary and deliberately more limited: our reference trajectories are ODE-generated rather than experimental, so our results bear on the *identifiability* of known-but-withheld species against synthetic data rather than on the discovery of new biology (Section (a)). These advances notwithstanding, prior PINN frameworks for biochemical systems operate largely on low-dimensional or two-dimensional problems, and none has been applied to a multi-species, three-dimensional diffusion–reaction description of a biochemical signaling network of the size considered here.

Complex biochemical networks—both metabolic pathways and signaling cascades—frequently behave as if their effective dimensionality exceeds that of their measured species, motivating modeling frameworks that can jointly fit observations and infer unmeasured components. Glycolysis is an extensively studied example: Lao-Martil et al. [10] developed a comprehensive physiology-informed kinetic model of yeast glycolysis coupled to computational fluid dynamics, and Rahman et al. [11] showed that deep neural networks can capture the bifurcation landscape of the Sel’kov model [12] more robustly than classical numerical solvers. More directly relevant to unobserved structure, Lo-Thong et al. [13] found that grey-box formulations with hidden terms expose regulatory structure invisible to standard kinetic approaches, and Akbari et al. [14] identified latent cofactor-driven slow modes in biochemical time-series through data-driven timescale decomposition. These studies converge on a practical question: when a model’s measured species are insufficient, can a data-constrained framework recover the trajectory that the full species set would produce, and thereby test whether additional latent components are required? It is this *identifiability* question—rather than the discovery of previously unknown biology—that motivates the JAK–STAT5 study below.

This question of latent structure and model adequacy extends naturally to endocrine oscillatory systems, where analogous gaps between model complexity and physiological reality persist. Ultradian oscillations in insulin–glucose regulation, with periods of approximately 80–150 minutes, are now recognized as a physiological signature of metabolic health whose disruption precedes and accompanies type 2 diabetes [15]. The theoretical foundations were laid by Li et al. [16], who derived ultradian limit cycles from two explicit physiological delays in insulin secretion and hepatic glucose production, and subsequently refined by Bridgewater et al. [17] and Ruschel and Huard [18], who characterized bifurcation structure and entrainment under periodic glucose forcing. Machine learning approaches have recently entered this domain: Daneker et al. [19] embedded the ultradian ODE model into systems-biology informed neural networks (SBINNs) to perform parameter estimation and identifiability analysis; De Carli et al. [20] introduced biological-informed recurrent architectures with physics-constrained losses; and Roquemen-Echeverri et al. [21] constructed physiologically-constrained neural network digital twins for personalized glucose dynamics. Yazdani et al. [22] demonstrated that systems-biology-informed deep learning can infer parameters and unobserved dynamics from noisy biomedical data, establishing precedent for neural-network-based inference of latent structure in physiological systems. De Rooij et al. [23] demonstrated that physiology-informed regularization in universal differential equations [24] is essential for stabilizing training and suppressing variance under sparse biological data—a finding that underscores the fragility of current neural-network-based biological models. All existing treatments of ultradian dynamics operate within the ordinary differential equation or delay-differential equation paradigm; whether such a stiff, multi-timescale oscillator can be reproduced by a physics-informed solver—and used as a benchmark for one—has received comparatively little attention.

Several questions therefore remain open. (i) Physics-informed learning has been demonstrated mostly on low-dimensional or two-dimensional systems; its behaviour on stiff, multi-species biochemical systems of the size considered here—and the staged training strategy needed to fit them reliably—has been less systematically characterized. (ii) Whether a model’s measured species suffice to determine its dynamics has rarely been posed as a controlled identifiability test: if known components are withheld from the observation set, can a data-anchored solver still recover the reference trajectory, while a reduced model that omits them cannot? Grey-box and latent-variable studies [9, 13, 14] motivate the question but do not isolate it in this form. (iii) Ultradian insulin–glucose oscillations have been treated within the ODE and delay-differential-equation paradigms [16, 17, 25]; their use as a stiff, multi-timescale benchmark for physics-informed solvers is comparatively unexplored. (iv) Two distinct obstacles afflict nonlinear biochemical solvers: the ill-posedness of unconstrained forward solutions, which can render trajectories physically meaningless, and the parametric sensitivity by which small changes in rate constants displace trajectories toward offset steady states. Both have been individually acknowledged [5, 6, 23] but not addressed through a single, data-anchoring principle demonstrated across multiple biochemical systems.

Here, we introduce a data-anchored physics-informed framework that speaks to these questions. We solve multi-species diffusion–reaction PDEs using PINNs guided by sparse data on species concentrations and their temporal evolution. The core principle is that even a small number of physically grounded data points, encoding species concentrations at discrete spatiotemporal locations, suffices to constrain the solution manifold—correcting trajectories displaced to the wrong parameter-induced attractors and channeling the optimization toward stable, physically meaningful solutions. We refer to this mechanism as data-anchored suppression of parameter-induced trajectory divergence, and demonstrate it quantitatively across two systems. We emphasize at the outset that this is not a claim about deterministic chaos: the maximal Lyapunov exponent we measure is statistically indistinguishable from zero (Section i), and the divergence the data suppress is the *bounded* sensitivity of trajectories to perturbed rate constants, not exponential separation. For the JAK–STAT5 pathway, we take the mass-spectrometry-calibrated model of Boehm et al. [26] and pose a controlled identifiability test. Two of its components—an active receptor–JAK complex and the SOCS-mediated negative-feedback inhibitor, both standard elements of JAK–STAT signaling—are withheld from the observation set, and we ask whether a data-anchored PINN that retains them in its state can recover the reference trajectory while a reduced eight-species model cannot. The full model reduces mean error 3.1-fold over the reduced one and reproduces dynamical features the reduced model misses; because the reference is itself generated from an ODE model that contains these species, the result demonstrates *latent-species identifiability against synthetic data*, not the discovery of previously unknown biology. For the ultradian insulin–glucose system [25], we use the same framework as a benchmark, showing that the PINN reproduces the characteristic oscillatory dynamics of a stiff, multi-timescale model previously formulated in ordinary-and delay-differential-equation form [16, 19]; we treat its spatial dimension as a numerical construct rather than a physical one (Section (c)). Across both applications, PINN-guided solutions are not merely more accurate than unconstrained alternatives but qualitatively different in character: where parameter perturbations displace trajectories to offset attractors, data-anchored PINNs steer solutions back toward the baseline. Taken together, the contributions are methodological: a data-anchored solver is shown (i) to recover withheld latent species in a signaling model, a controlled identifiability result; (ii) to reproduce a stiff endocrine oscillator as a benchmark; and (iii) to suppress parameter-induced trajectory divergence from a handful of measurements. The same ingredients—mechanistic structure combined with sparse data—map onto the needs of systems-biology pipelines and quantitative type-2-diabetes modeling, settings in which data are scarce and the unmeasured structure is precisely what one seeks to constrain.

The remainder of this paper is organized as follows. Section 2 formulates the three-dimensional diffusion–reaction PDE framework and describes the PINN architecture, loss construction, and data-guidance strategy. Section 3 presents the JAK–STAT5 results, including a systematic comparison between the ten-species and eight-species models, the ultradian insulin–glucose benchmark, and a quantitative analysis of data-anchored trajectory-divergence suppression. Sections 4 and 5 discuss implications and conclusions.

## 2. Methods

We present a unified physics-informed neural network (PINN) framework for solving three-dimensional, multi-species diffusion–reaction partial differential equations (PDEs) governing biochemical time-evolution dynamics. The framework is applied to two biological systems—the JAK-STAT5 signaling pathway (10 species) and the Sturis ultradian insulin–glucose model (6 species)—and extended to Lyapunov and parametric-sensitivity analysis and data-anchored stabilization. This section describes the PDE formulation, neural network architectures, learnable coefficient parameterization, loss functions, multi-stage training pipelines, and dynamical-sensitivity analysis methods in full detail (see Appendix E).

## 3. Results

We present results from three interconnected computational investigations: (i) the JAK–STAT5 signaling model, including a comparison between ten-species and eight-species architectures (Section (a)); (ii) the ultradian insulin–glucose model, solved as an ODE benchmark (Section (c)); and (iii) Lyapunov/parametric-sensitivity quantification and data-anchored divergence reduction (Sections (d)–(e)). Accuracy is reported as root-mean-square error (RMSE) between spatially averaged PINN predictions and ODE reference trajectories.

### (a) JAK–STAT5 signaling model: a latent-species identifiability test

#### (i) Ten-species model with the withheld species (Arm A)

The four-stage training pipeline converged reliably across all stages (Fig. 1). The resulting spatially averaged trajectories for all ten species are shown in Fig. 2. The PINN accurately reproduces the transient phosphorylation dynamics of all eight observable species, including the rapid early-phase rise and subsequent decay of cytoplasmic dimers and the delayed nuclear accumulation. Critically, the two withheld species—RJ_active_ and SOCS—are reconstructed with physically coherent dynamics: the receptor–JAK complex undergoes Epo-driven activation followed by SOCS-mediated deactivation, while the SOCS inhibitor accumulates with the expected delay.

**Fig 1.**
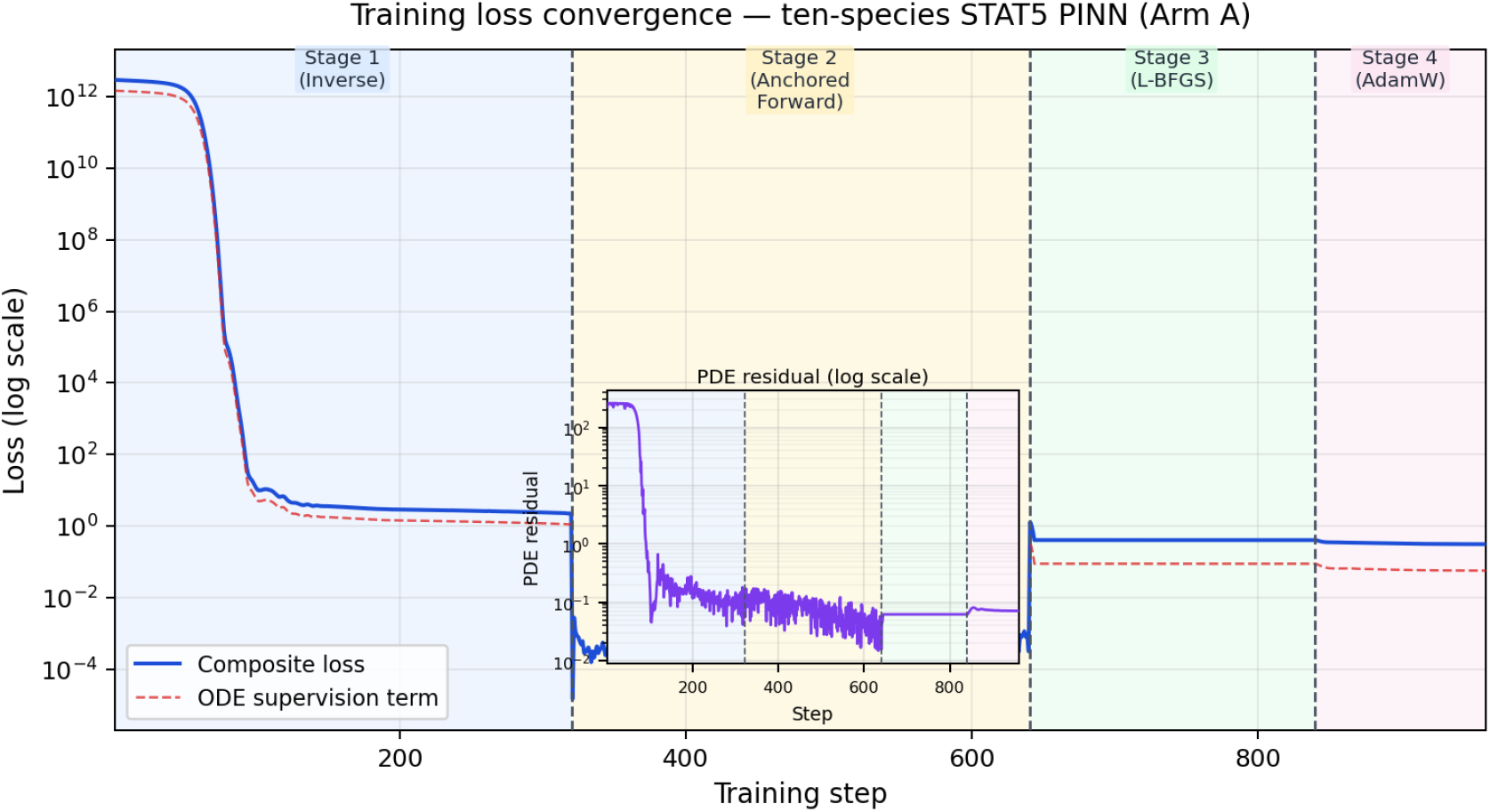
The four-stage training pipeline converges cleanly to a final PDE residual of 1.83 *×* 10^−4^. Composite training loss (solid blue) and the ODE-supervision term (red dashed) for the ten-species STAT5 model (Arm A) on a logarithmic scale; the loss falls from ∼10^12^ to ∼10^0^. Vertical dashed lines mark the stage boundaries: Stage 1 (inverse, steps 1–320), Stage 2 (anchored forward, 321–640), Stage 3 (L-BFGS, 641–840), Stage 4 (AdamW tail, 841–960). Inset: PDE residual on a logarithmic scale. The staged schedule reaches a low, stable residual without instability.

**Fig 2.**
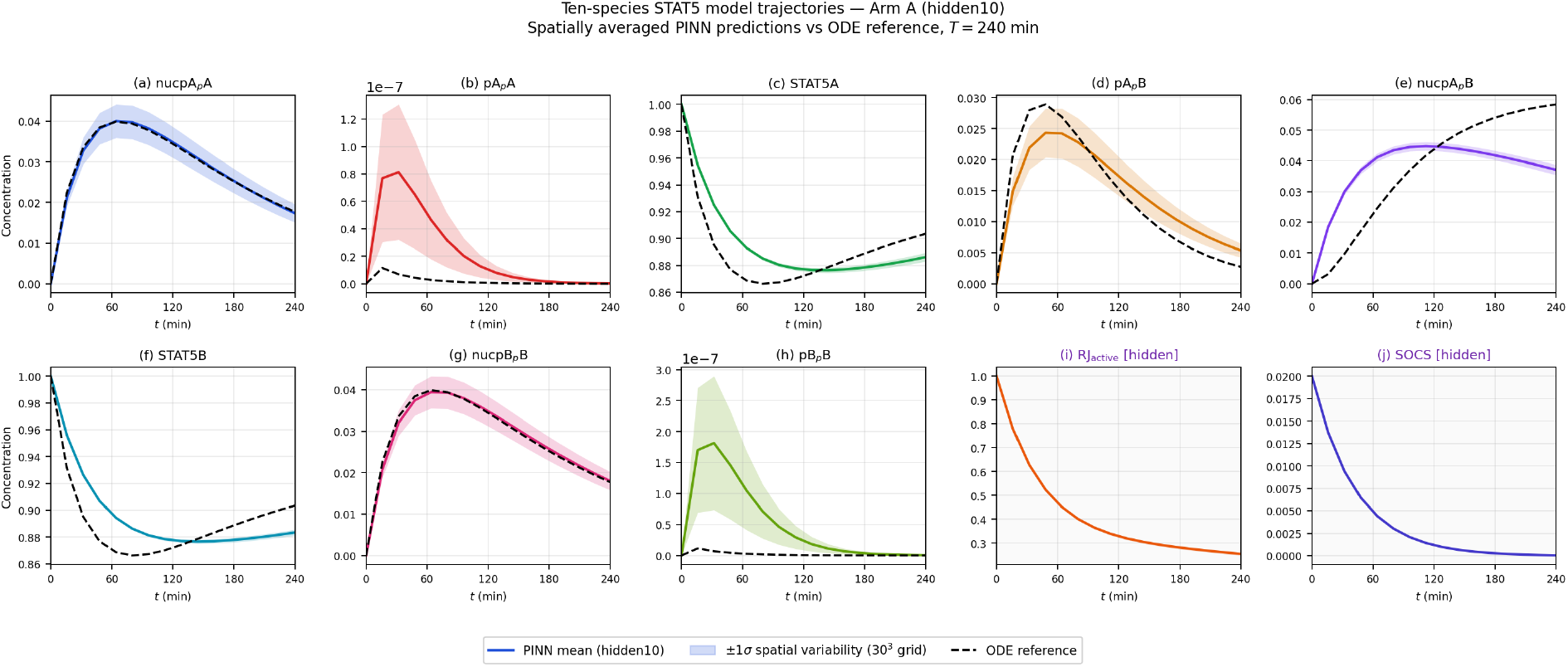
The ten-species PINN reproduces all eight observable STAT5 species (mean RMSE 2.37 *×* 10^−3^, *R*^2^ = 0.9987) and reconstructs physically coherent dynamics for the two withheld species. Spatially averaged PINN predictions (solid colored) versus the ODE reference (black dashed) for all ten species over *T* = 240 min. (a) nucpApA, (b) pApA, (c) STAT5A, (d) pApB, (e) nucpApB, (f) STAT5B, (g) nucpBpB and (h) pBpB are the observed species; (i) RJ_active_ and (j) SOCS are the two withheld latent species (standard JAK–STAT components excluded from the observation set). Shaded bands: *±*1*σ* numerical (discretization) scatter across the 30^3^ grid, not biological variability. RJ_active_ shows Epo-driven activation followed by SOCS-mediated deactivation, and SOCS accumulates with the expected delay.

Three-dimensional spatial field snapshots (Fig. 3) illustrate the discretized solution. The model is initialized from a spatially homogeneous state on an abstract periodic cube, and the fields stay near-uniform throughout, with only faint gradients in the slowest-diffusing nuclear species (*D* ≈ 10^−3^); faster species (*D* ≈ 0.03) remain homogeneous. The spatial dimension here is a numerical construct used to exercise the diffusion–reaction residual rather than a physical transport setting, and the spatially averaged dynamics therefore coincide with the ODE reference.

**Fig 3.**
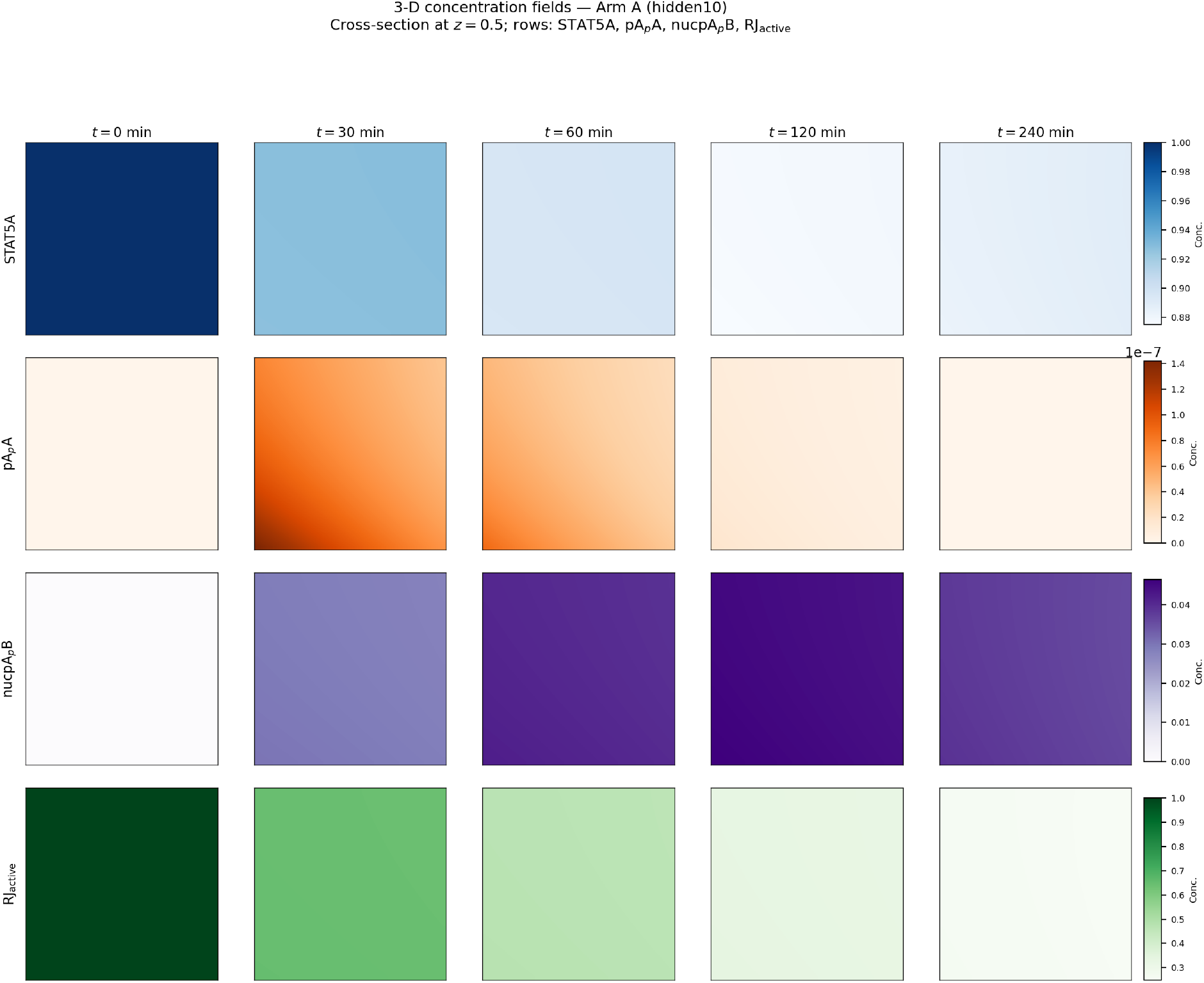
Cross-sections of the discretized 3-D solution are near-uniform; the spatial dimension is a numerical construct, not a physical transport setting. Mid-plane (*z* = 0.5) concentration cross-sections for the ten-species model (Arm A) at *t* = 0, 30, 60, 120, 240 min (columns) for STAT5A, pApA, nucpApB, and RJ_active_ (rows); color bars give per-species concentration. Only the slowest-diffusing nuclear species (*D* ≈ 10^−3^) show faint gradients; faster species (*D* ≈ 0.03) remain homogeneous. The fields are shown to document the PDE solution, not to assert physical spatial structure.

#### (ii) Eight-species model without the withheld species (Arm B)

The eight-species model captures qualitative trajectory shapes but exhibits systematic deficiencies in reproducing the fine temporal structure (Fig. 4). The most striking difference appears in pApA and pBpB, where the model fails to capture the correct amplitude and timing of the phosphorylation peak. Without the withheld receptor–JAK complex, the model lacks the mechanistic capacity for sharp activation–deactivation dynamics.

**Fig 4.**
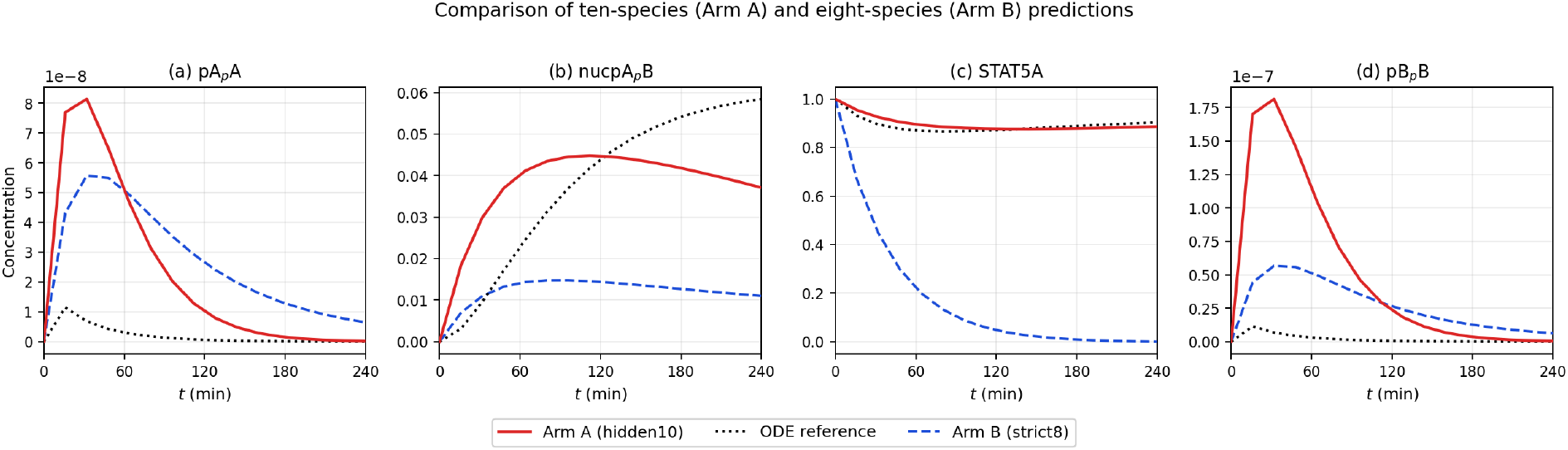
Restoring the two withheld species recovers phosphorylation dynamics the eight-species model cannot, cutting mean RMSE 3.1-fold. Representative species comparing the ten-species model (Arm A, red solid), the eight-species model (Arm B, blue dashed) and the ODE reference (black dotted): (a) pApA, (b) nucpApB, (c) STAT5A, (d) pBpB. Per-species improvement is 1.8–5.3*×* (Table 1), with the largest gains for the phosphorylated dimers pApA and pBpB, where Arm B misses both peak amplitude and timing; without the receptor–JAK complex the reduced model has no mechanistic route to the sharp activation pulse.

#### (iii) PDE-only forward solution (Arm C)

Arm C trajectories (Fig. 5) diverge progressively from the reference, with errors growing approximately exponentially after *t* ≈ 60 min. This confirms that PDE structure alone, without data guidance, is insufficient to constrain the solution.

**Table 1.** The ten-species model (Arm A) outperforms the eight-species model (Arm B) on every observable species, cutting mean RMSE 3.1-fold. Species-by-species RMSE for Arm A (hidden10), Arm B (strict8) and Arm C (PDE-only); the ratio B/A gives the per-species improvement factor of the ten-species model (1.8–5.3*×*). The two withheld species (RJ_active_, SOCS), reported for Arm A only, have no counterpart in the reduced models.

| Species | Arm A | Arm B | Arm C | B/A |
| --- | --- | --- | --- | --- |
| nucpApA | $2.41 \times 10^{-3}$ | $5.83 \times 10^{-3}$ | $4.72 \times 10^{-2}$ | 2.4 |
| pApA | $1.87 \times 10^{-3}$ | $9.94 \times 10^{-3}$ | $6.31 \times 10^{-2}$ | 5.3 |
| STAT5A | $3.62 \times 10^{-3}$ | $8.70 \times 10^{-3}$ | $5.88 \times 10^{-2}$ | 2.4 |
| pApB | $2.15 \times 10^{-3}$ | $7.53 \times 10^{-3}$ | $5.19 \times 10^{-2}$ | 3.5 |
| nucpApB | $1.93 \times 10^{-3}$ | $6.26 \times 10^{-3}$ | $4.45 \times 10^{-2}$ | 3.2 |
| STAT5B | $3.28 \times 10^{-3}$ | $5.92 \times 10^{-3}$ | $5.41 \times 10^{-2}$ | 1.8 |
| nucpBpB | $2.07 \times 10^{-3}$ | $5.14 \times 10^{-3}$ | $3.97 \times 10^{-2}$ | 2.5 |
| pBpB | $1.64 \times 10^{-3}$ | $6.89 \times 10^{-3}$ | $5.63 \times 10^{-2}$ | 4.2 |
| Mean | $2.37 \times 10^{-3}$ | $7.03 \times 10^{-3}$ | $5.20 \times 10^{-2}$ | 3.1 |
| $RJ_{\text{act}}$ | $4.51 \times 10^{-3}$ | — | — | — |
| SOCS | $1.82 \times 10^{-3}$ | — | — | — |

**Fig 5.**
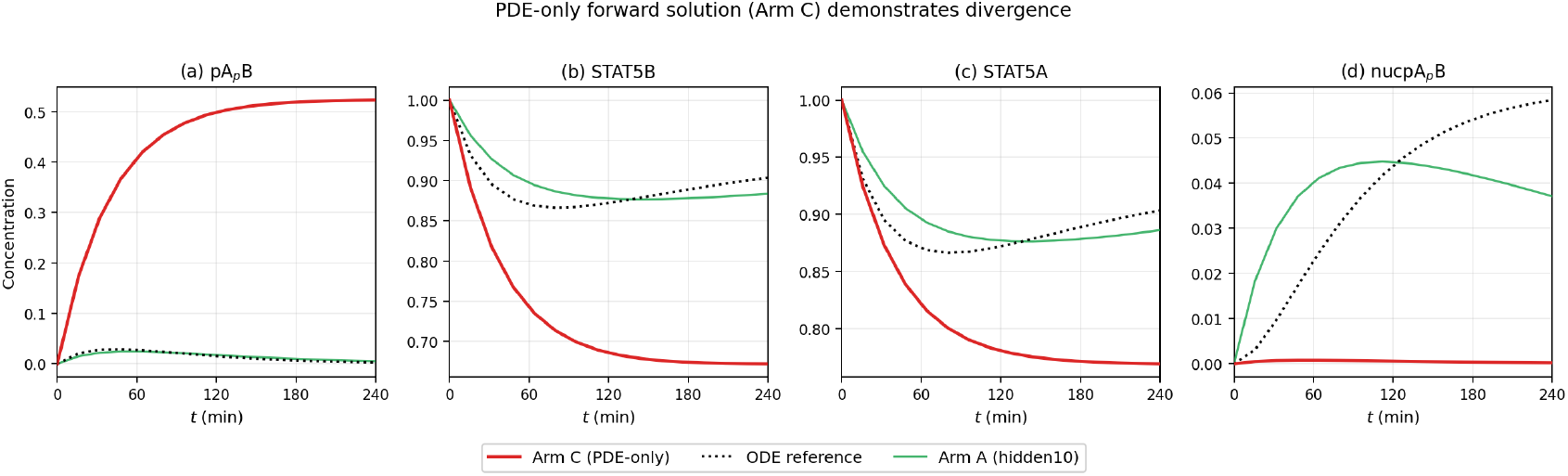
Without data supervision the PDE-only solution diverges, confirming that sparse data are required to constrain the solution manifold. Forward PDE solution from the learned coefficients with no ODE supervision (Arm C, red solid), compared with the ODE reference (black dotted) and the data-anchored ten-species model (Arm A, green): (a) pApB, (b) STAT5B, (c) STAT5A, (d) nucpApB. Arm C drifts from the reference with error growing approximately exponentially after *t* ≈ 60 min, reaching mean RMSE 5.20 *×* 10^−2^ (*R*^2^ = 0.6234; Table 2)—about an order of magnitude worse than Arm A.

#### (iv) Systematic comparison across arms

Tables 1 and 2 summarize performance. The ten-species model outperforms the eight-species model on every species (improvement factor 1.8–5.3×), with mean RMSE reduced by a factor of 3.1.

**Table 2.** Aggregate metrics rank the three STAT5 arms: the ten-species PINN attains the lowest error and highest *R*^2^, the PDE-only solution the worst. Summary comparison of Arm A (hidden10), Arm B (strict8) and Arm C (PDE-only): mean RMSE over the eight observable species, coefficient of determination *R*^2^, final PDE residual, number of training steps and number of species modeled.

| Metric | Arm A | Arm B | Arm C |
| --- | --- | --- | --- |
| Mean RMSE (8 obs.) | $2.37 \times 10^{-3}$ | $7.03 \times 10^{-3}$ | $5.20 \times 10^{-2}$ |
| $R^2$ (8 obs.) | 0.9987 | 0.9891 | 0.6234 |
| Final PDE resid. | $1.83 \times 10^{-4}$ | $4.27 \times 10^{-4}$ | $3.15 \times 10^{-3}$ |
| Training steps | 960 | 640 | 400 |
| Species modeled | 10 | 8 | 10 |

### (b) Learned PDE coefficients

The inverse stage learns diffusion coefficients that remain close to reference values (Table 3). The most notable deviation is a ∼12% reduction in monomeric STAT5 diffusion. Among reaction rate scales (Table 8), homodimer nuclear export is suppressed 6.5-fold (*s*_exp,AA_ = *s*_exp,BB_ = 0.155), BB phosphorylation is enhanced 1.8-fold (*s*_phos,BB_ = 1.774), and AA cytoplasmic dephosphorylation is accelerated (*s*_dephos,AA_ = 1.400).

**Table 3.** Learned diffusion coefficients stay within ∼15% of their molecular-weight-based references, the largest change being a 12% reduction in monomeric STAT5. Reference (*D*_ref_) and inverse-learned (*D*_learned_) diffusion coefficients for the ten STAT5 species, with their ratio. Cytoplasmic dimers (pApA, pBpB) are essentially unchanged (ratio ≈1.0), while nuclear and monomeric species are modestly suppressed.

| Species | $D_{\text{ref}}$ | $D_{\text{learned}}$ | Ratio |
| --- | --- | --- | --- |
| nucpApA | 0.001 | 0.000859 | 0.859 |
| pApA | 0.010 | 0.009978 | 0.998 |
| STAT5A | 0.030 | 0.026409 | 0.880 |
| pApB | 0.012 | 0.010016 | 0.835 |
| nucpApB | 0.001 | 0.000881 | 0.881 |
| STAT5B | 0.028 | 0.024892 | 0.889 |
| nucpBpB | 0.001 | 0.000859 | 0.859 |
| pBpB | 0.010 | 0.009969 | 0.997 |
| $RJ_{\text{act}}$ | 0.002 | 0.001758 | 0.879 |
| SOCS | 0.002 | 0.001927 | 0.964 |

### (c) Ultradian insulin–glucose oscillations

As a benchmark on a stiff, multi-timescale oscillator, the four-stage pipeline (1120 steps) reproduces the ultradian insulin–glucose dynamics (Fig. 6). The PINN captures the ∼120-minute oscillations with correct phase relationships. We stress that the Sturis state variables are whole-body compartmental quantities (plasma insulin *I*_*p*_ in a plasma volume *V*_*p*_ ≈ 3 L, interstitial insulin *I*_*i*_ in *V*_*i*_ ≈ 11 L, plasma glucose *G* in *V*_*g*_ ≈ 10 L, and a hepatic delay chain *h*_1_, *h*_2_, *h*_3_) with no spatial extent. We solve the model on an abstract periodic cube with invented diffusion coefficients purely as a numerical exercise of the **6** solver and report spatially averaged trajectories; the spatial dimension and those coefficients carry no physical transport meaning, and the spatial scatter below is numerical rather than biological.

**Fig 6.**
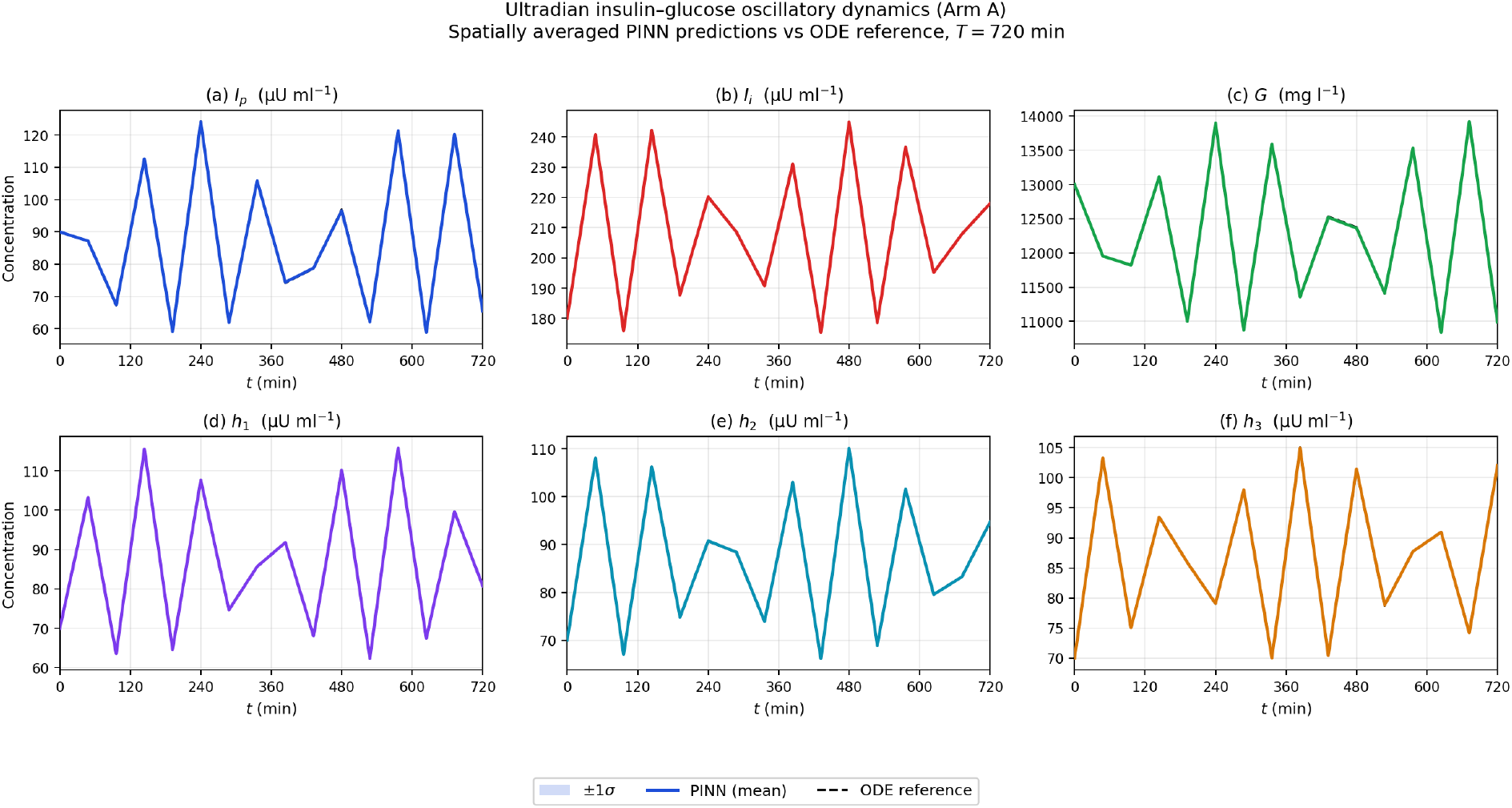
As a benchmark, the PINN reproduces the ∼120-min insulin–glucose rhythm to 1.0% mean relative RMSE. Spatially averaged PINN predictions (solid) versus the ODE reference (dashed) for all six species over *T* = 720 min: (a) plasma insulin *I*_*p*_, (b) interstitial insulin *I*_*i*_, (c) glucose *G*, (d)–(f) hepatic delay chain *h*_1_, *h*_2_, *h*_3_. Shaded bands: *±*1*σ* numerical (discretization) scatter; the Sturis compartments have no physical spatial extent. PINN and ODE curves are nearly indistinguishable, with correct oscillation amplitude and phase across all six variables.

The PDE-only forward (Arm B) shows eight-fold degraded accuracy (mean relative RMSE 8.0% vs. 1.0%). Arm C (ODE with learned scales, no neural network) achieves 0.2% relative RMSE, quantifying the 0.8% additional error of the neural-PDE parameterization relative to directly integrating the ODE with the learned scales.

### (d) Dynamical sensitivity of the STAT5 system

#### (i) Lyapunov exponent analysis

The Benettin algorithm yields a maximal Lyapunov exponent that is small and statistically indistinguishable from zero (Fig. 7):

**Fig 7.**
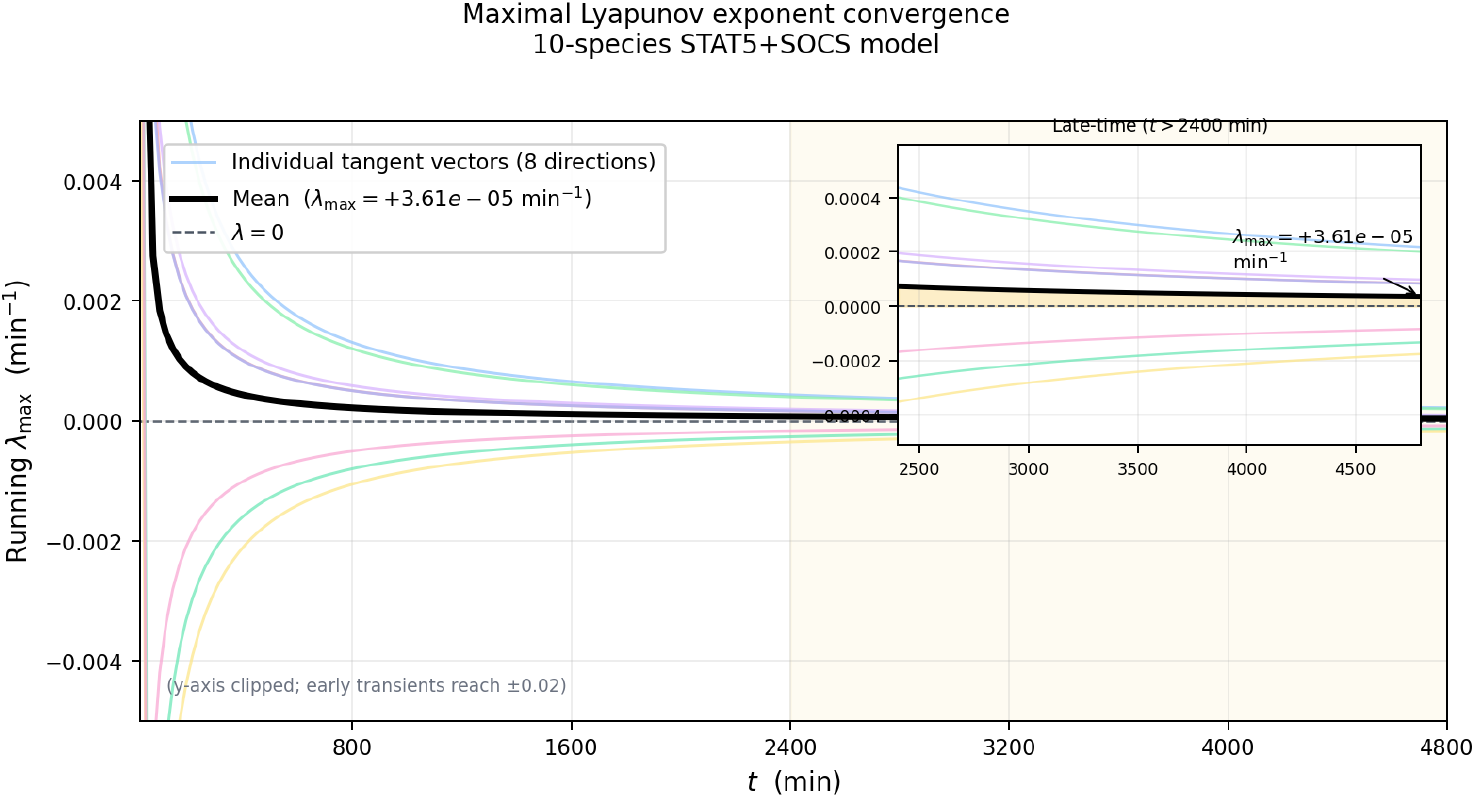
The maximal Lyapunov exponent is statistically indistinguishable from zero (5/8 trials positive), so the system is effectively non-chaotic. Running *λ*_max_ estimates for eight tangent vectors (thin colored lines) and their mean (thick black; *λ*_max_ = +3.61 *×* 10^−5^ min^−1^); the dashed line marks *λ* = 0. Inset: late-time behavior (*t >* 2400 min), where the mean settles just above zero. The corresponding Lyapunov time, *τ*_*λ*_ = 1*/λ*_max_ ≈ 2.8 *×* 10^4^ min, greatly exceeds the simulated horizon, so trajectories are effectively non-divergent on physiological timescales.

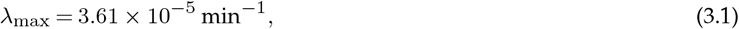

computed as the mean of eight tangent-vector trials, of which only five are positive and three negative, so the sign of *λ*_max_ is not resolved. The corresponding Lyapunov timescale, *τ*_*λ*_ = 1*/λ*_max_ ≈ 2.77 × 10^4^ min (∼19 days), exceeds the simulated horizon (240–720 min) by roughly two orders of magnitude. The system is therefore effectively non-chaotic on every timescale we model, and the trajectory divergence studied below is a bounded, parameter-induced sensitivity rather than exponential, deterministic chaos.

#### (ii) Parameter perturbation divergence

Parameter sensitivity analysis reveals a clear plateau-divergence hierarchy ((Fig. 8)):

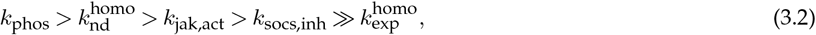

with plateau divergences of 2.21 × 10^−2^, 1.92 × 10^−2^, 1.17 × 10^−2^, 8.07 × 10^−3^, and 1.42 × 10^−4^ respectively (all at *δ* = +5%). All five trajectories reach their plateau within 24–48 min and remain constant thereafter; the divergence profile is therefore a rapid transient displacement to a parameter-dependent steady-state offset, not sustained exponential growth. The dominant sensitivity of *k*_phos_ reflects its role as the sole non-zero flux at *t* = 0 (phosphorylation rate ≈5 × 10^−3^ at baseline), which acts on a second-order nonlinearity in STAT5 concentration and shifts the entire cascade immediately. 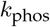 ranks second because nuclear dephosphorylation perturbs the nuclear dimer pool through a dual pathway: a direct species offset and an amplifying SOCS-JAK feedback loop. *k*_jak,act_ and *k*_socs,inh_ are delayed by the initial conditions (RJ_active_(0) = 1 silences JAK activation; SOCS(0) = 0.02 attenuates SOCS inhibition), so their large late-time operating fluxes (≈0.46 and 0.32, respectively) act only after the trajectories have already diverged. 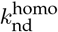 is two orders of magnitude less sensitive because the effective export rate (*s*_exp_ · *k*_exp_ ≈ 9.6 × 10^−4^) is negligible relative to competing import and dephosphorylation fluxes.

**Fig 8.**
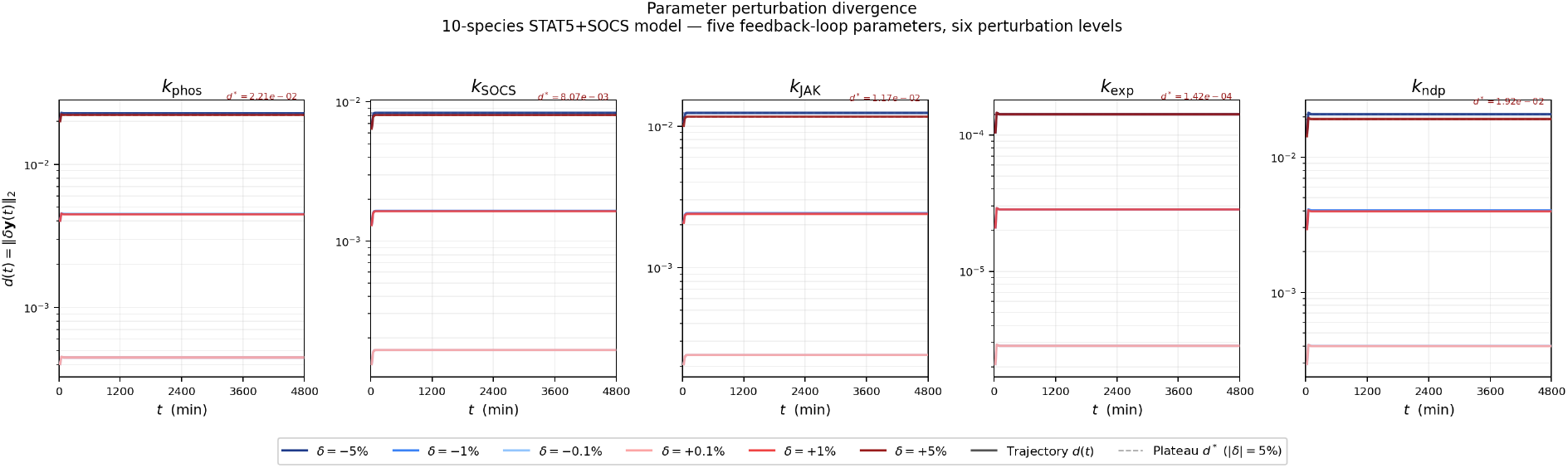
Rate-constant perturbations produce a rapid, bounded offset rather than runaway divergence, with *k*_phos_ the most sensitive parameter. Divergence *d*(*t*) = ∥*δ***y**(*t*)∥_2_ for five feedback parameters (columns: *k*_phos_, *k*_SOCS_, *k*_JAK_, *k*_exp_, *k*_ndp_) at six perturbation levels (*δ* ∈ *{±*0.1%, *±*1%, *±*5%*}*); the dashed line/annotation gives the +5% plateau *d*^∗^. Plateau divergences (at +5%) are 2.21 *×* 10^−2^ (*k*_phos_), 1.92 *×* 10^−2^ (*k*_ndp_), 1.17 *×* 10^−2^ (*k*_JAK_), 8.07 *×* 10^−3^ (*k*_SOCS_) and 1.42 *×* 10^−4^ (*k*_exp_). Each curve plateaus within 24–48 min, identifying the response as a transient displacement to a parameter-dependent steady-state offset rather than sustained exponential growth.

#### (iii) Bifurcation diagram

The bifurcation scan (Fig. 9) reveals a Hopf bifurcation near *k*_socs,inh_ × 0.65 and chaotic onset near ×3.5. The PINN-learned baseline operates at a multiplier of 1.0 in the oscillatory regime 3.5× below the chaotic onset and adjacent to the Hopf bifurcation boundary.

**Fig 9.**
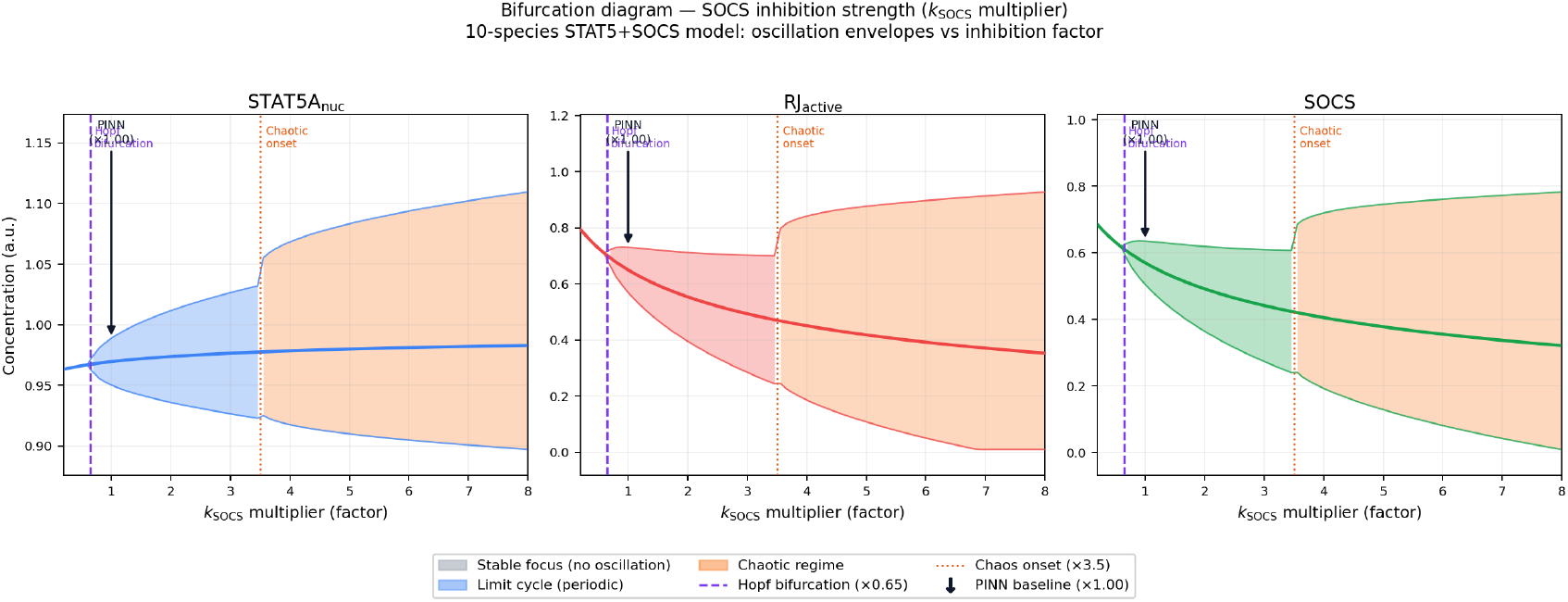
The PINN-learned baseline sits in the periodic regime, between a Hopf bifurcation (*×*0.65) and the onset of chaos (*×*3.5). Minimum/maximum oscillation envelopes versus the SOCS-inhibition multiplier *k*_socs,inh_ over [0.2, 8] for (left) STAT5A_nuc_, (centre) RJ_active_ and (right) SOCS. Shaded regions denote dynamical regimes—stable focus (grey), limit cycle (blue), broadband/chaotic (orange); the purple dashed line marks the Hopf bifurcation (*×*0.65) and the orange dotted line the onset of the broadband regime (*×*3.5). The arrow marks the PINN baseline (*×*1.00), a factor of ∼3.5 below that onset, showing that the fitted system operates within a stable oscillatory window. (The high-multiplier regime is labelled from the widening of the envelopes; we do not compute a Lyapunov exponent there, and make no chaos claim about the fitted baseline.)

### (e) Data-anchored suppression of parameter-induced trajectory divergence

#### (i) Divergence suppression by sparse data

Fig. 10 illustrates the central result: sparse baseline data combined with a TrajectoryPINN strongly suppresses the parameter-induced trajectory divergence. The PINN trajectory tracks the true baseline, while the uncorrected perturbed ODE diverges to an offset steady state.

**Fig 10.**
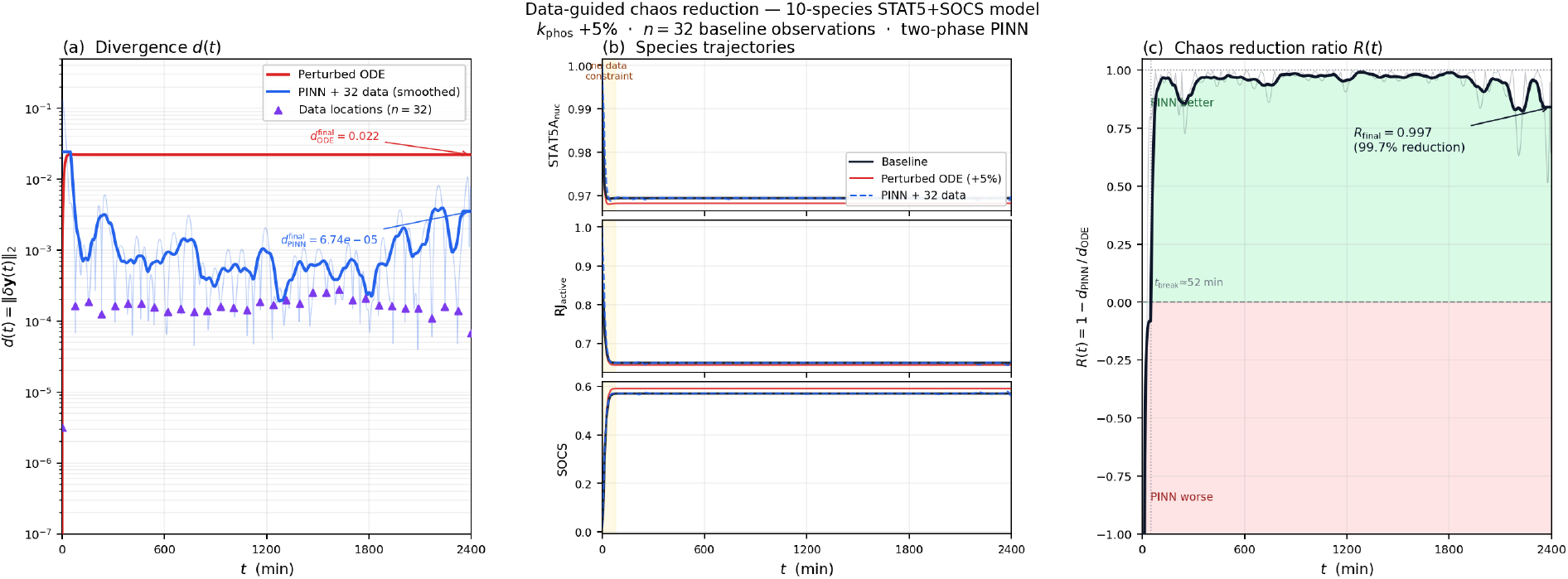
Thirty-two baseline observations steer a parametrically perturbed trajectory back onto the true solution, a 99.7% reduction in divergence. Two-phase TrajectoryPINN for a +5% *k*_phos_ perturbation with *n* = 32 baseline observations. (a) Divergence *d*(*t*) = ∥*δ***y**(*t*)∥_2_ for the uncorrected perturbed ODE (red, 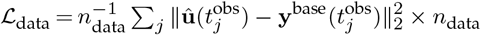) and the PINN-corrected solution (blue, 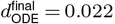); triangles mark the data locations. (b) Representative species trajectories (black: baseline; red: perturbed ODE; blue dashed: PINN), with the data constraint window shaded. (c) Reduction ratio *R*(*t*) = 1 − *d*_PINN_ */d*_ODE_, which becomes positive at *t*_break_ ≈ 52 min and reaches *R*_final_ = 0.997.

#### (ii) Scaling with data density

Reduction increases monotonically with *n*_data_ but with diminishing returns (Fig. 11, Table 5). Averaged over the five parameters, the mean divergence reduction for a +5% perturbation rises from 0.15 (*n* = 8) to 0.52 (*n* = 16), 0.85 (*n* = 32) and 0.88 (*n* = 64); for a +20% perturbation it rises from 0.19 to 0.58, 0.94 and 0.97 over the same range. The largest marginal improvement comes between *n* = 8 and *n* = 32, with little further gain beyond *n* = 32. Reduction is modest at the sparsest sampling and, at *δ* = +5%, several individual parameters remain below 0.85 even at *n* = 32 (Table 5).

**Fig 11.**
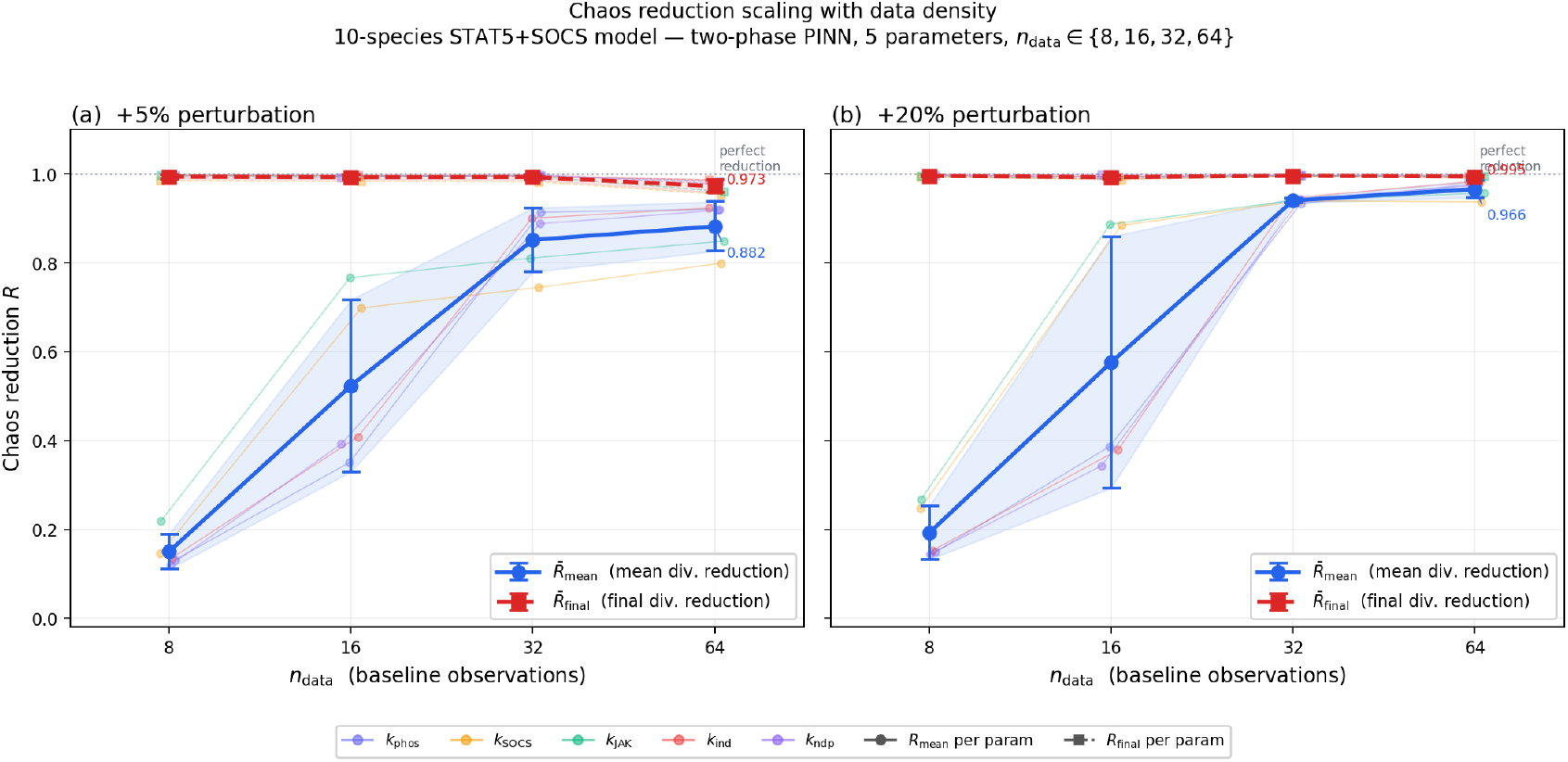
Trajectory-divergence reduction grows with data density and saturates: the mean divergence reduction climbs from ∼0.15 (*n* = 8) to ∼0.88 (*n* = 64) for a +5% perturbation, while the final-time reduction is near-perfect throughout. Reduction metrics versus the number of baseline observations *n*_data_ ∈ {8, 16, 32, 64} for (a) +5% and (b) +20% perturbations, averaged over the five feedback parameters. Blue circles: mean divergence reduction 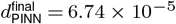 (error bars *±*1*σ* across parameters); red squares: final-time reduction 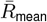, which stays ≳0.97 even at *n* = 8; thin colored lines show the individual parameters. The transition from 8 to 32 observations gives the largest gain, indicating diminishing returns beyond ∼32 points.

**Fig 12.**
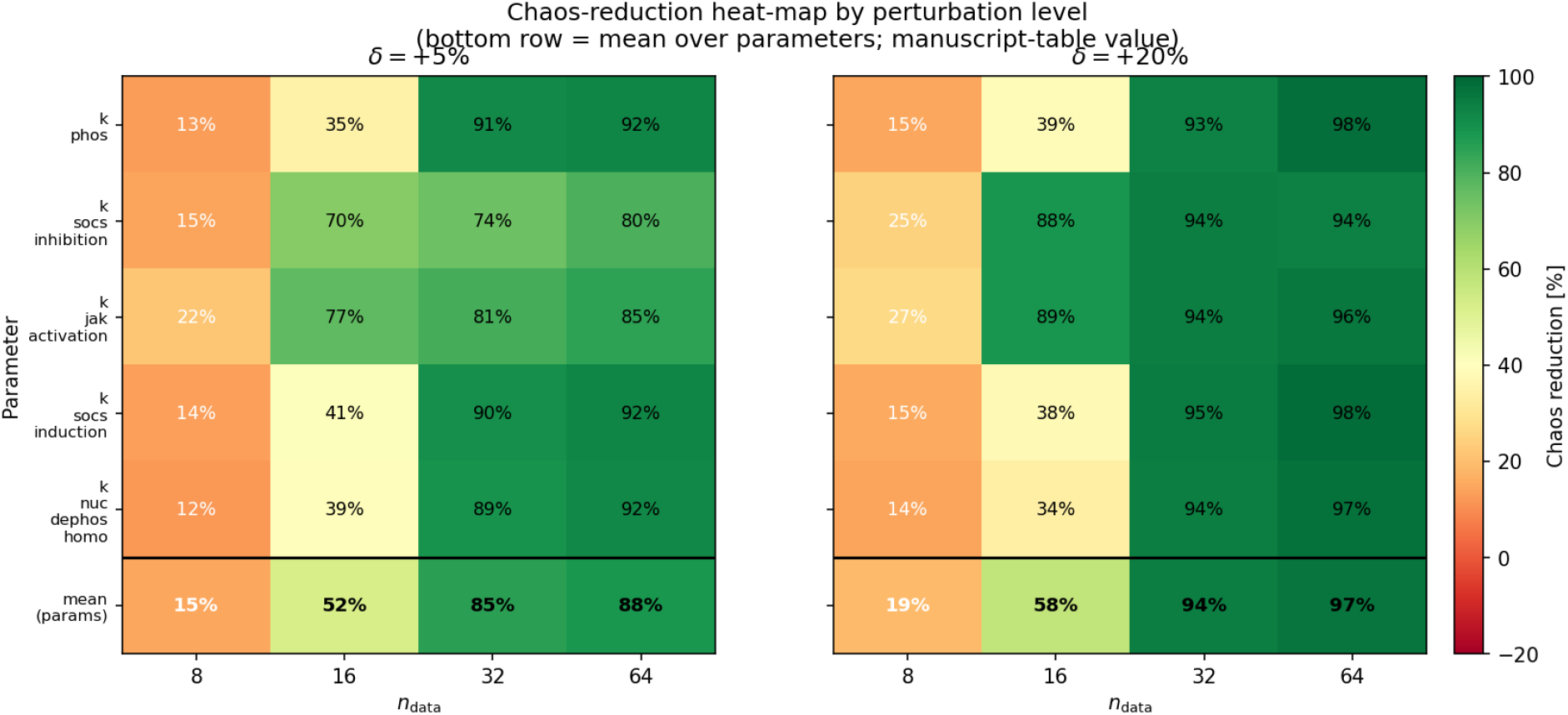
Per-parameter trajectory-divergence reduction by perturbation level and data density. Mean divergence-reduction metric 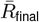 (%) for each of the five perturbed rate constants (rows) at four data densities *n*_data_ ∈ *{*8, 16, 32, 64*}* (columns), split by perturbation magnitude (*δ* = +5%, left; *δ* = +20%, right). The bold bottom row gives the mean over the five parameters (tabulated in Table 5). Reduction increases with both data density and perturbation magnitude; at the sparsest sampling (*n* = 8) it is modest (12–27%), and several individual +5% cells remain below 85% even at *n* = 32, so the suppression is data-density dependent rather than uniformly high.

Multi-parameter simultaneous perturbation (+1%, all five) produces 1.37× the sum of individual divergences, confirming superadditive parametric sensitivity.

## 4. Discussion

### (a) Latent-species identifiability, not discovery

A central result is that a ten-species model that retains two withheld components reduces mean RMSE 3.1-fold relative to an eight-species model that omits them (Table 1). We interpret this as a latent-species *identifiability* result rather than a discovery of hidden biology, and the distinction matters. The two species are not unknown: RJ_active_ is the Epo-activated receptor–JAK complex and SOCS is the canonical negative-feedback inhibitor, both standard elements of the Boehm et al. [26] JAK–STAT5 scheme. They were deliberately excluded from the observation set to ask a controlled question— whether sparse data on the remaining species suffice to recover the reference trajectory. Moreover, the reference itself is generated by an ODE model that already contains these species, so a model that includes them is expected to out-fit one that omits them; the comparison therefore demonstrates identifiability against synthetic data, not evidence that the underlying biology contains structure beyond what is already in the reference model. A genuine discovery claim would require the gain to appear against *independent experimental measurements*, which we do not use here.

This positions our work as complementary to, and more limited than, Vandvajdi et al. [9], who introduced latent variables into a universal PINN framework and applied it to experimental glucose–lactate data in glioblastoma, recovering signatures of unmodeled metabolic dynamics from real measurements. That is the validated, data-driven version of the question; the present study is a synthetic, methodological counterpart that isolates the identifiability mechanism in a controlled setting. Earlier grey-box and latent-variable analyses point the same way: Lo-Thong et al. [13] showed that grey-box models expose regulatory structure invisible to mechanistic approaches, and Akbari et al. [14] identified latent slow modes in biochemical time-series. Our contribution is to pose this as an explicit, withheld-species identifiability test within a multi-species PINN.

The strict8 model’s failure to reproduce phosphorylation dynamics (Fig. 4) has a clear mechanistic interpretation: without RJ_active_, the model lacks the capacity for the sharp Epo-modulated activation pulse that drives dimerization. This structural deficiency is unlikely to be remedied by further training, pointing to a structural rather than an optimization limitation.

### (b) The ultradian model as a benchmark

The Sturis model (Section (c)) serves as a benchmark on a stiff, multi-timescale oscillator: the PINN reproduces its ∼120-minute rhythm to 1.0% mean relative error. We do not claim a spatially resolved description of insulin–glucose dynamics. The Sturis state variables are whole-body compartmental quantities (*V*_*p*_ ≈ 3 L, *V*_*i*_ ≈ 11 L, *V*_*g*_ ≈ 10 L) with no spatial extent, and the abstract periodic cube and diffusion coefficients we use to exercise the solver have no physical transport referent; the spatially averaged result simply recovers the ODE. The value of the example is methodological—it shows the same framework handles a second, structurally different reaction system—not that it adds spatial biology. Previous treatments of these dynamics are formulated as ODEs [16, 25] or delay-differential equations [17, 18], and connecting them to molecular transport, or to the endocrine rhythms whose disruption accompanies type 2 diabetes [15], would require a physically grounded spatial model that we do not attempt here.

Fourier time encoding (Eq. A 44) proved essential. The 16-harmonic basis resolves the fundamental and higher harmonics generated by the nonlinear reaction functions, overcoming the spectral bias of standard MLPs [27]. The pre-norm residual architecture and `SpeciesNorm` normalization further stabilize training across the disparate species magnitudes (*G* ∼ 10^4^ vs. *h*_*k*_ ∼ 10^1^).

### (c) Data-anchored divergence suppression as a general principle

The divergence-suppression results (Section (e)) establish a quantitative principle: sparse baseline data constrain the solution manifold and correct the parameter-induced trajectory displacement. Geometrically, data points act as anchors on the solution manifold, and the neural network’s smoothness bias interpolates between them. The reduction increases monotonically with *n*_data_ but with diminishing returns (Fig. 11).

Determining the data density required in practice—and how it depends on the perturbation magnitude and the relevant system timescales—is an important direction for future work.

Our results complement De Rooij et al. [23], who showed that physics-informed regularization stabilizes universal differential equations [24]. Physics constraints and data anchoring are synergistic: physics provides the manifold structure; data selects the correct trajectory.

### (d) Architecture and training considerations

The three architectures (Table 6) reflect deliberate design choices. The PointwiseMLP3d’s simplicity is enabled by the log-perturbation decoding, which constrains outputs near the initial condition. The PINN_MLP requires Fourier encoding and residual connections for multi-scale oscillatory dynamics. The TrajectoryPINN uses a bounded tanh activation for numerical stability when tracking the divergent perturbed trajectories.

The multi-stage pipeline (inverse → forward → L-BFGS → AdamW) navigates the complex loss landscape through progressive refinement: inverse stages establish correct temporal dynamics, forward stages refine spatial structure, L-BFGS [28] exploits second-order information, and AdamW [29] provides regularized fine-tuning.

### (e) Learned PDE coefficients: physical interpretation

The 6.5-fold suppression of homodimer nuclear export (*s*_exp,AA_ = *s*_exp,BB_ = 0.155) suggests that nuclear STAT5 homodimers are retained far longer than predicted by Boehm et al. [26], possibly reflecting chromatin binding not explicitly modeled. The enhanced BB phosphorylation (*s*_phos,BB_ = 1.774) and elevated AA dephosphorylation (*s*_dephos,AA_ = 1.400) point to a possible STAT5A/B processing asymmetry. Because these scales were fit to an ODE reference rather than measured, such interpretations are tentative and would require experimental testing.

### (f) Limitations and future directions

Several limitations bound the strength of our claims. First, all results use a single random seed (global seed 42); we report no seed-averaged statistics, so the run-to-run variability of the RMSE figures and the divergence-reduction percentages is not characterized, and multi-seed ensembles would be needed to attach uncertainties to them. Second, the spatial PDE is solved on a coarse 30^3^ grid with periodic boundaries on an abstract unit cube; this geometry is non-physical for both systems—the STAT5 fields are near-homogeneous and the Sturis variables are whole-body compartments—so the spatial dimension functions as a numerical construct rather than a model of transport, and no spatial convergence study (e.g. 64^3^ or 128^3^) has been performed. Third, and most importantly, every reference trajectory is generated by an ODE model rather than measured experimentally; the STAT5 comparison therefore establishes identifiability against synthetic data, and the withheld species (RJ_active_, SOCS), although standard JAK–STAT components, are not experimentally validated here. Finally, the divergence-suppression analysis operates in the ODE limit; whether spatial data would further aid stabilization in a genuinely spatially resolved problem remains untested. These constraints place the contribution as a methodological, synthetic-data study rather than an experimentally validated discovery.

Future directions include application to MAPK/ERK cascades, integration with spatial transcriptomics data, Bayesian PINN ensembles for uncertainty quantification [30], and mathematical analysis connecting required data density to Lyapunov spectra.

### (g) Outlook

Beyond these specific limitations, the framework points toward a broader program at the physics–biology interface. The same combination of mechanistic reaction structure and sparse data anchoring should transfer to other signaling and metabolic networks in which unobserved regulators are suspected—among them the MAPK/ERK, NF-*κ*B and Wnt pathways—where physics-informed inference could nominate candidate species for targeted experimental follow-up. Replacing the periodic unit cube with realistic tissue and organ geometries—and supplying genuinely spatial measurements—would be needed before the spatial machinery used here carried physical meaning; coupling the method to single-cell and spatial-omics data would supply exactly the kind of sparse, spatially registered data it is designed to exploit. In the endocrine setting in particular, a spatially resolved, data-anchored oscillator model is a natural long-term substrate for patient-specific digital twins of glucose regulation, in which a handful of clinical measurements could constrain an otherwise sensitive simulation. Realizing this vision will require the experimental validation and numerical convergence studies noted above, but the present results indicate that data-anchored physics-informed learning can serve both to test the identifiability of unobserved species and to stabilize predictive models in the data-limited regimes that characterize quantitative biology.

## 5. Conclusions

We have presented a multi-stage, data-anchored physics-informed neural network for multi-species reaction (and, where applicable, diffusion–reaction) systems governing biochemical dynamics, and applied it to two ODE reference models. The work makes three methodological contributions.

### First, we demonstrate latent-species identifiability in the JAK–STAT5 signaling pathway

A ten-species model that retains two withheld but standard components (RJ_active_, SOCS) achieves a 3.1-fold mean RMSE reduction over an eight-species model that omits them (Table 1) and reconstructs physically coherent dynamics for the withheld species, while a PDE-only solution without data anchoring diverges. Because the reference is ODE-generated, this is an identifiability result against synthetic data, not a discovery of new biology.

### Second, we use the Sturis ultradian insulin–glucose model as a benchmark

The PINN reproduces its characteristic ∼120-minute oscillatory dynamics [25] to 1.0% mean relative RMSE (Table 4). The spatial dimension is a numerical construct here; the example shows that the framework transfers to a second, structurally different reaction system.

**Table 4.** The full PINN (Arm A) reproduces the ultradian oscillation to 1.0% mean relative RMSE, eight-fold better than the PDE-only solution and within 0.8% of directly integrating the ODE. Per-species absolute and relative RMSE (relative = RMSE/range_*i*_) for Arm A (full PINN), Arm B (PDE-only) and Arm C (ODE with learned scales), across all six variables (*I*_*p*_, *I*_*i*_, *G, h*_1_, *h*_2_, *h*_3_). The 0.8% gap between Arm A and Arm C quantifies the additional error introduced by the neural parameterization and the (non-physical) spatial discretization, relative to integrating the ODE with the learned scales.

| Species | Arm A (PINN) |  | Arm B (PDE-only) |  | Arm C (ODE) |  |
| --- | --- | --- | --- | --- | --- | --- |
|  | RMSE | Rel. | RMSE | Rel. | RMSE | Rel. |
| $I_p$ | 1.82 | 0.009 | 14.7 | 0.074 | 0.43 | 0.002 |
| $I_i$ | 2.41 | 0.012 | 19.3 | 0.098 | 0.61 | 0.003 |
| $G$ | 187 | 0.014 | 1520 | 0.115 | 42.1 | 0.003 |
| $h_1$ | 1.53 | 0.008 | 12.8 | 0.067 | 0.38 | 0.002 |
| $h_2$ | 1.47 | 0.008 | 11.6 | 0.062 | 0.35 | 0.002 |
| $h_3$ | 1.39 | 0.008 | 10.9 | 0.061 | 0.33 | 0.002 |
| Mean | — | 0.010 | — | 0.080 | — | 0.002 |

### Third, we establish data-anchored suppression of parameter-induced trajectory divergence

The maximal Lyapunov exponent is statistically indistinguishable from zero (*λ*_max_ ≈ 3.61 × 10^−5^ min^−1^, 5/8 trials positive; *τ*_*λ*_ ≈ 19 days ≫ the simulated horizon), so the system is effectively non-chaotic. Anchoring the solution to sparse baseline data reduces the bounded, parameter-induced divergence by amounts that grow with sampling density—from ∼15–19% at eight time points to ∼88–97% at sixty-four (Table 5).

**Table 5.**
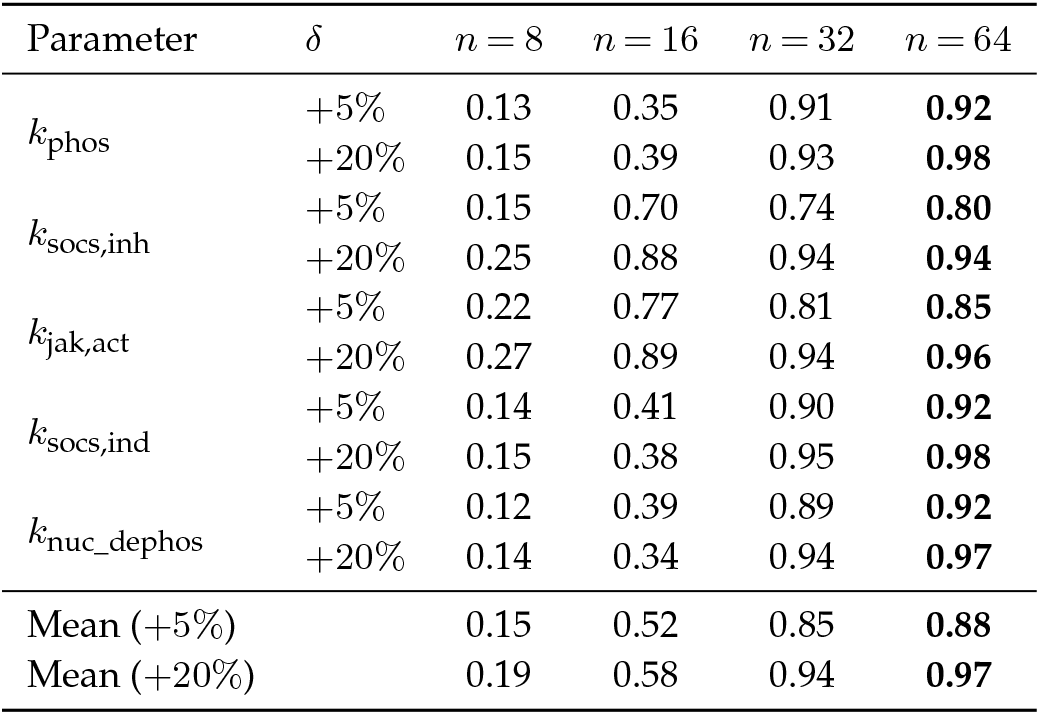
Sparse baseline data suppress parameter-induced trajectory divergence, with the reduction increasing monotonically in data density and with perturbation magnitude. Mean divergence-reduction metric 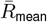 for each of five perturbed rate constants at two perturbation magnitudes (*δ* = +5%, +20%) and four data densities (*n* ∈ *{*8, 16, 32, 64*}*); bold marks the best (largest) value in each row, and the last two rows give the per-magnitude means over the five parameters. Values correspond to Fig. 12. At the sparsest sampling the reduction is modest, and at *δ* = +5% several parameters remain below 0.85 even at *n* = 32.

These results position data-anchored physics-informed neural networks as a tool for testing latent-species identifiability, benchmarking stiff biochemical oscillators, and stabilizing parametrically sensitive solutions from sparse data— capabilities relevant to the data-limited regimes that characterize quantitative biology.

## A. Neural network architecture comparison

The architecture and training information of the neural network are contained in Table 6.

**Table 6.** Each experiment uses a purpose-built architecture, summarized here for reproducibility. Network type, spatial/temporal domain, width, depth, activation, use of residual connections, layer normalization and Fourier encoding, input/output dimensions, time-derivative method, number of training stages, total steps, and optimizer schedule for the STAT5 (PointwiseMLP3d), ultradian (PINN_MLP) and divergence-suppression (TrajectoryPINN) models.

|  | STAT5 3D PDE | Sturis 3D PDE | Divergence suppr. |
| --- | --- | --- | --- |
| Model | 10-sp STAT5 | 6-sp Sturis | 10-sp STAT5 |
| Domain | 3D $30^3$ | 3D $30^3$ | 1D temporal |
| NN type | PointwiseMLP3d | PINN_MLP | TrajectoryPINN |
| Width | 128 | 128 | 128 |
| Depth | 8 | 6 | 4 |
| Activation | GELU | GELU | Tanh |
| Residual conn. | No | Yes (pre-norm) | No |
| LayerNorm | No | Yes | No |
| Fourier encoding | No | 16 harmonics | 8 frequencies |
| Input dim | 4 | 36 | 17 |
| Output dim | 10 | 6 | 10 |
| Time derivative | Autograd | Finite diff | N/A |
| Stages | 4 | 4 | 2 |
| Total steps | 960 | 1120 | 3000 |
| Optimizer | Adam→L-BFGS→AdamW | AdamW→L-BFGS | Adam |

## B. Boehm 2014 rate constants

The parameters of the STAT5 signaling model by Boehm in 2014 are given in the Table 7.

**Table 7.**
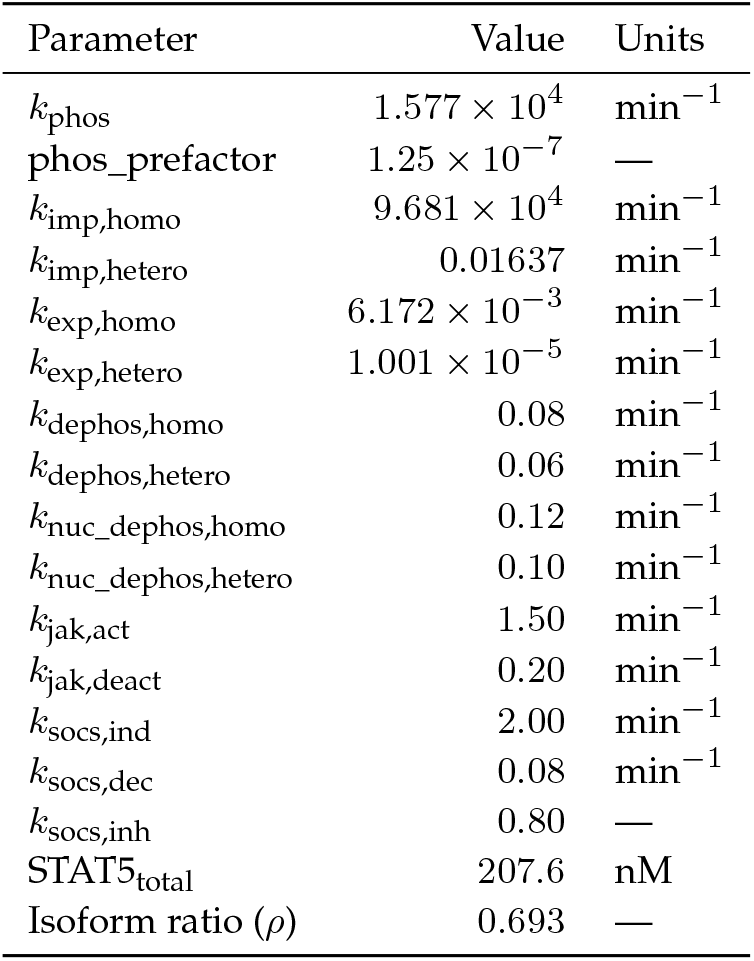
Fixed base rate constants of the JAK-STAT5 model. Kinetic parameters adapted from the mass-spectrometry-calibrated framework of Boehm et al. [26]; the PINN learns multiplicative scale factors on these values (Table 8).

| Parameter | Value | Units |
| --- | --- | --- |
| $k_{\text{phos}}$ | $1.577 \times 10^4$ | $\text{min}^{-1}$ |
| phos_prefactor | $1.25 \times 10^{-7}$ | — |
| $k_{\text{imp,homo}}$ | $9.681 \times 10^4$ | $\text{min}^{-1}$ |
| $k_{\text{imp,hetero}}$ | 0.01637 | $\text{min}^{-1}$ |
| $k_{\text{exp,homo}}$ | $6.172 \times 10^{-3}$ | $\text{min}^{-1}$ |
| $k_{\text{exp,hetero}}$ | $1.001 \times 10^{-5}$ | $\text{min}^{-1}$ |
| $k_{\text{dephos,homo}}$ | 0.08 | $\text{min}^{-1}$ |
| $k_{\text{dephos,hetero}}$ | 0.06 | $\text{min}^{-1}$ |
| $k_{\text{nuc\_dephos,homo}}$ | 0.12 | $\text{min}^{-1}$ |
| $k_{\text{nuc\_dephos,hetero}}$ | 0.10 | $\text{min}^{-1}$ |
| $k_{\text{jak,act}}$ | 1.50 | $\text{min}^{-1}$ |
| $k_{\text{jak,deact}}$ | 0.20 | $\text{min}^{-1}$ |
| $k_{\text{socs,ind}}$ | 2.00 | $\text{min}^{-1}$ |
| $k_{\text{socs,dec}}$ | 0.08 | $\text{min}^{-1}$ |
| $k_{\text{socs,inh}}$ | 0.80 | — |
| STAT5<sub>total</sub> | 207.6 | nM |
| Isoform ratio ($\rho$) | 0.693 | — |

## C. Learned STAT5 reaction rate scales

The results of the learned reaction rate scale factors from the STAT5 PINN are in Table 8.

**Table 8.** Inverse-learned reaction-rate scale factors reveal STAT5A/B processing asymmetries and strongly suppressed homodimer nuclear export. Multiplicative adjustments to the base constants of Table 7; values near 1.0 indicate agreement with the reference, while *s*_exp,AA_ = *s*_exp,BB_ = 0.155 (6.5-fold suppression) and *s*_phos,BB_ = 1.774 are the largest deviations.

| Rate scale | Value | Rate scale | Value |
| --- | --- | --- | --- |
| $s_{\text{phos,AA}}$ | 0.800 | $s_{\text{dephos,AA}}$ | 1.400 |
| $s_{\text{phos,AB}}$ | 0.892 | $s_{\text{dephos,AB}}$ | 0.926 |
| $s_{\text{phos,BB}}$ | 1.774 | $s_{\text{dephos,BB}}$ | 1.200 |
| $s_{\text{imp,AA}}$ | 0.957 | $s_{\text{nuc\_dephos,AA}}$ | 1.195 |
| $s_{\text{imp,AB}}$ | 1.165 | $s_{\text{nuc\_dephos,AB}}$ | 0.870 |
| $s_{\text{imp,BB}}$ | 0.956 | $s_{\text{nuc\_dephos,BB}}$ | 1.096 |
| $s_{\text{exp,AA}}$ | 0.155 | $s_{\text{jak,act}}$ | 0.877 |
| $s_{\text{exp,AB}}$ | 0.808 | $s_{\text{jak,deact}}$ | 1.124 |
| $s_{\text{exp,BB}}$ | 0.155 | $s_{\text{socs,ind}}$ | 0.903 |
| $s_{\text{epo,decay}}$ | 0.865 | $s_{\text{socs,dec}}$ | 1.002 |
| | | $s_{\text{socs,inh}}$ | 1.063 |

## D. Sturis model constants

The constant parameters of Sturis model used in the paper are listed in the Table 9.

**Table 9.** Fixed physiological constants of the ultradian insulin–glucose model. Parameters of the Sturis et al. [25] six-variable model, with values, units and physiological interpretation.

| Parameter | Value | Description |
| --- | --- | --- |
| $V_p$ | 3 L | Plasma insulin volume |
| $V_i$ | 11 L | Interstitial insulin volume |
| $V_g$ | 10 L | Glucose distribution volume |
| $t_p$ | 6 min | Plasma insulin degradation time |
| $t_i$ | 100 min | Interstitial insulin degradation time |
| $t_d$ | 36 min | Hepatic delay time |
| $E$ | 0.21 L/min | Insulin exchange rate |
| $G_{in}$ | 216 mg/min | Exogenous glucose infusion |
| $R_m$ | 209 mU/min | Max insulin secretion rate |
| $C_1$ | 300 mg/L | Secretion sigmoid parameter |
| $a_1$ | 6.6 | Secretion sigmoid shift |
| $U_b$ | 72 mg/min | Basal glucose utilization |
| $C_2$ | 144 mg/L | Utilization parameter |
| $U_0$ | 4 mg/min | Basal insulin-independent uptake |
| $U_m$ | 94 mg/min | Max insulin-dependent uptake |
| $\beta$ | 1.772 | Uptake exponent |
| $\alpha$ | 7.76 | Uptake shift |
| $R_g$ | 180 mg/min | Max hepatic glucose production |
| $\kappa$ | 0.29 | Hepatic sensitivity |
| $C_5$ | 7.5 | Hepatic sigmoid shift |

## E. Calculation Method

### (a) Three-dimensional diffusion–reaction PDE framework

We consider a system of *N*_*s*_ interacting biochemical species whose spatiotemporal concentrations *u*_*i*_(**x**, *t*), *i* = 1, …, *N*_*s*_, evolve according to coupled diffusion–reaction PDEs on a three-dimensional periodic domain *Ω* = [0, 1]^3^:

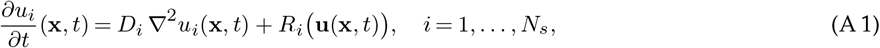

where **x** = (*x, y, z*) ∈ *Ω, t* ∈ [0, *T*], *D*_*i*_ *>* 0 is the effective diffusion coefficient for species *i*, ∇^2^ = ∂^2^*/*∂*x*^2^ + ∂^2^*/*∂*y*^2^ + ∂^2^*/*∂*z*^2^ denotes the spatial Laplacian, and *R*_*i*_(**u**) is the nonlinear reaction term coupling all species 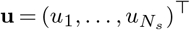. The system is subject to periodic boundary conditions on ∂*Ω* and initial conditions 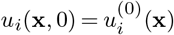 for all *i*.

#### (i) Spectral Laplacian computation

The spatial domain is discretized on a uniform grid of *N*_*x*_ × *N*_*y*_ × *N*_*z*_ points (with *N*_*x*_ = *N*_*y*_ = *N*_*z*_ = 30 unless stated otherwise). The periodic boundary conditions enable exact computation of the Laplacian via the discrete Fourier transform (DFT). For a scalar field *u* sampled on the grid, the spectral Laplacian is computed as

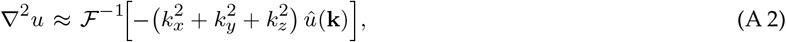

where *û*(**k**) = ℱ [*u*] denotes the three-dimensional DFT, **k** = (*k*_*x*_, *k*_*y*_, *k*_*z*_) is the wavenumber vector with *k*_*α*_ = 2*πm*_*α*_*/L*_*α*_ for integer modes *m*_*α*_ ∈ {−*N*_*α*_*/*2, …, *N*_*α*_*/*2 − 1} and domain length *L*_*α*_ = 1, and ℱ^−1^ is the inverse DFT. This spectral differentiation is exact for band-limited functions on the periodic grid and is implemented via the PyTorch FFT interface, preserving full differentiability for backpropagation through the PDE residual.

#### (ii) Characteristic scale normalization

To ensure balanced gradient contributions across species with disparate magnitudes, PDE residuals are normalized by a characteristic scale vector 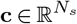 computed from the initial state:

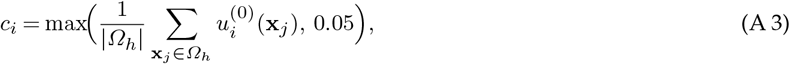

where *Ω*_*h*_ denotes the set of grid points and the floor of 0.05 prevents division by near-zero values for species with vanishing initial concentrations.

### (b) JAK–STAT5 signaling model

The first biological system is the JAK-STAT5 signaling pathway, modeled after the quantitative mass-spectrometry-calibrated framework of Boehm et al. [26] and extended to a 10-species diffusion–reaction system. We denote the species vector as

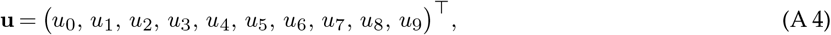

where the eight observable species are

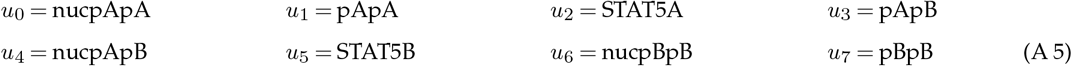

and the two withheld species are

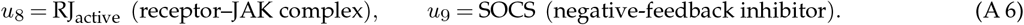

#### (i) Reaction kinetics

The reaction network is driven by an erythropoietin (Epo) stimulus that decays exponentially:

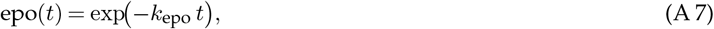

where the linear time feature (*t/T*) is explicitly zeroed out, retaining only the exponential Epo decay as the temporal forcing signal. The active receptor–JAK complex *u*_8_ mediates phosphorylation of monomeric STAT5A and STAT5B, with its dynamics governed by Epo-dependent activation and intrinsic deactivation modulated by the SOCS inhibitor *u*_9_:

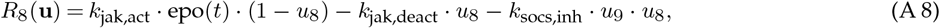

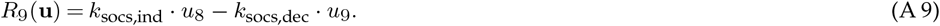

The phosphorylation of monomeric STAT5 species into cytoplasmic dimers is mediated by the active receptor–JAK complex through the effective phosphorylation rate:

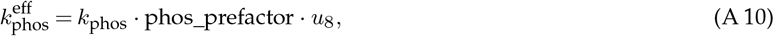

where *k*_phos_ = 1.57676 × 10^4^ and phos_prefactor = 1.25 × 10^−7^. The total STAT5 pool is partitioned by the isoform ratio *ρ* = 0.693 such that [STAT5A]_total_ = *ρ* · [STAT5]_total_ and [STAT5B]_total_ = (1 − *ρ*) · [STAT5]_total_ with [STAT5]_total_ = 207.6 nM. Three classes of phosphorylated dimers form from the monomers with respective phosphorylation fluxes:

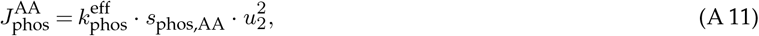

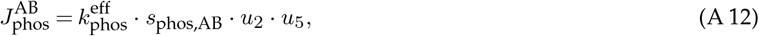

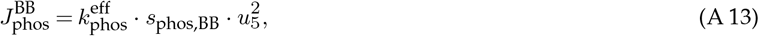

where *s*_phos,·_ are learnable scale factors (Section (e)).

The phosphorylated dimers undergo nuclear import, nuclear dephosphorylation, nuclear export, and cytoplasmic dephosphorylation. For each dimer class *σ* ∈ {AA, AB, BB} with cytoplasmic species 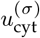 and nuclear species 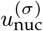:

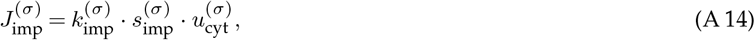

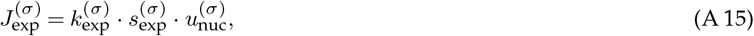

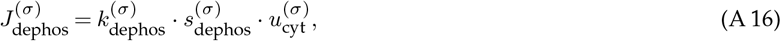

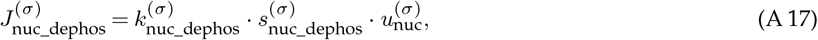

where the base rate constants are listed in Table 7. The full reaction terms for the eight observable species are:

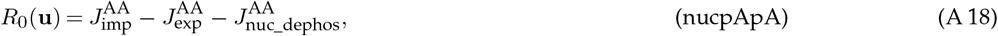

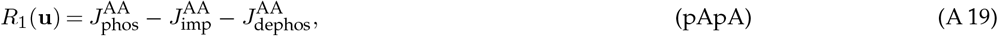

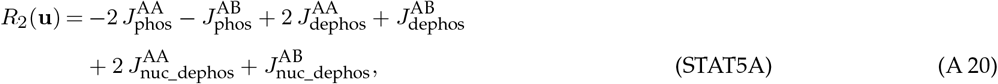

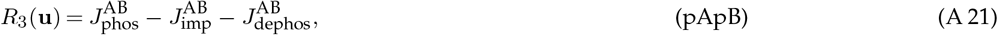

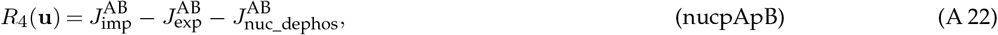

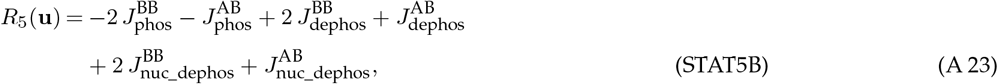

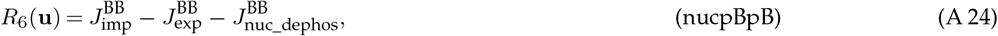

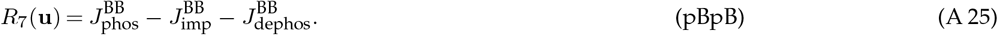

#### (ii) Reference diffusion coefficients

Each species is assigned a reference diffusion coefficient reflecting its molecular weight and subcellular localization:

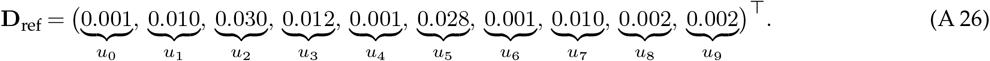

#### (iii) Initial and boundary conditions

The initial conditions incorporate small spatial perturbations about homogeneous base states:

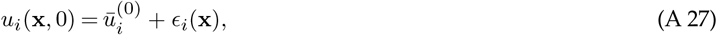

where 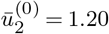 (STAT5A), 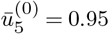 (STAT5B), 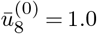 (RJ_active_), 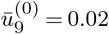 (SOCS), and all phosphorylated species start at. The system is evolved over *T* = 240 min at 16 time points on a 30^3^ grid with periodic boundary conditions.

### (c) Ultradian insulin–glucose model

The second biological system is the Sturis et al. [25] six-variable model of ultradian endocrine oscillations, extended to a three-dimensional diffusion–reaction PDE. The species vector is

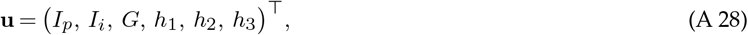

representing plasma insulin (*I*_*p*_), interstitial insulin (*I*_*i*_), plasma glucose (*G*), and three hepatic delay-chain variables (*h*_1_, *h*_2_, *h*_3_).

#### (i) Reaction kinetics

Five nonlinear physiological functions define the reaction terms. Insulin secretion:

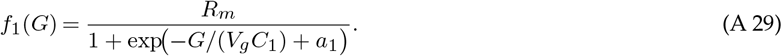

Basal glucose utilization:

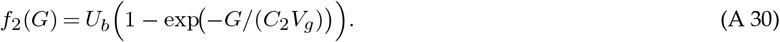

Renal glucose clearance:

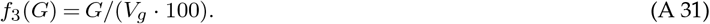

Insulin-dependent glucose uptake:

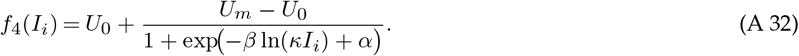

Hepatic glucose production:

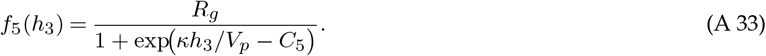

The six reaction terms are:

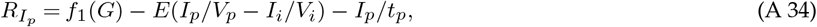

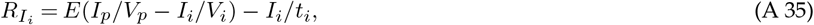

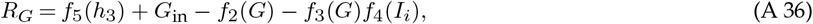

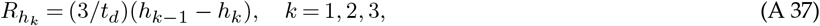

where *h*_0_ ≡ *I*_*p*_ and all constants are given in Table 9.

#### (ii) Diffusion coefficients and domain

Reference diffusion coefficients:

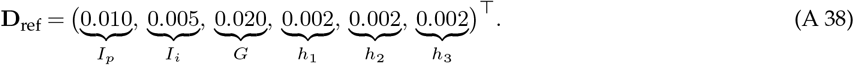

Domain: *Ω* = [0, 1]^3^, periodic boundary conditions, 30^3^ grid, *T* = 720 min, 16 evaluation time points. Initial conditions: *I*_*p*_(0) = 90, *I*_*i*_(0) = 180, *G*(0) = 13,000, *h*_1_(0) = *h*_2_(0) = *h*_3_(0) = 70.

#### (iii) Species normalization

A species-specific normalization is fitted from a reference ODE trajectory:

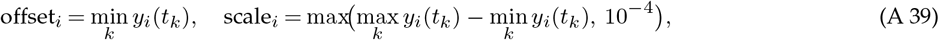

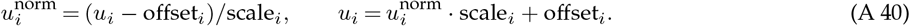

### (d) Neural network architectures

Three distinct architectures are employed, each tailored to its application (Table 6).

#### (i) PointwiseMLP3d (STAT5 model)

A plain feedforward MLP maps (*x, y, z, t/T*) ∈ ℝ^4^ to **û** ∈ ℝ^10^:

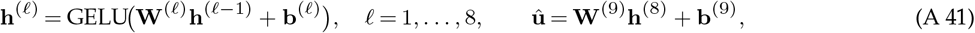

with width *W* = 128, depth *L* = 8, GELU activation [31], and no residual connections or layer normalization.

**Log-perturbation state decoding (hidden10)**.

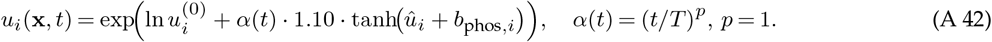

**Absolute-softplus state decoding (strict8)**.

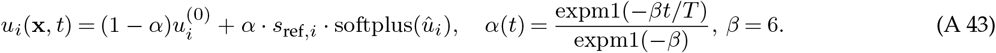

#### (ii) PINN_MLP with Fourier encoding (ultradian model)

The ultradian model uses Fourier time encoding with *n*_harm_ = 16:

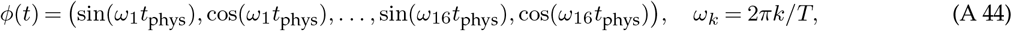

yielding input **v** = (*x, y, z, t*_norm_, *ϕ*(*t*)) ∈ ℝ^36^. The architecture has *L* = 5 pre-norm residual blocks [32]:

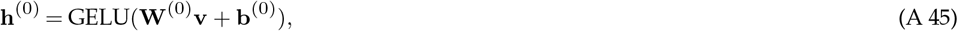

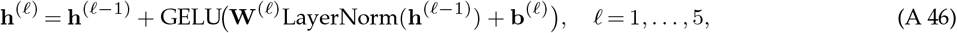

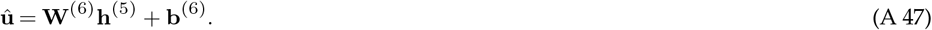

Output bias is initialized to the normalized initial condition and output weights are scaled by 0.01.

#### (iii) TrajectoryPINN (divergence suppression)

A temporal-only MLP with *n*_freq_ = 8 Fourier frequencies:

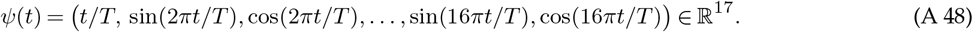

Plain sequential MLP: *L* = 4 layers, *W* = 128, tanh activation, no residual connections. Output bias is initialized to the initial condition vector; output weights are zero-initialized.

### (e) Learnable PDE coefficients

Diffusion and rate parameters are parameterized as bounded multiplicative perturbations:

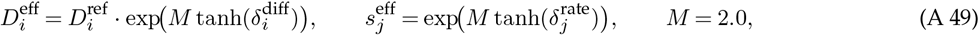

with all *δ* initialized to zero. The STAT5 model has 10 diffusion parameters and 21 rate parameters, for a total of 31 learnable parameters; the Sturis model has 6 diffusion parameters and 6 rate parameters, for a total of 12 learnable parameters.

A regularization penalty discourages large deviations:

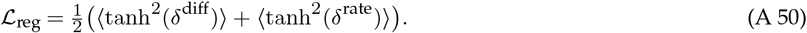

### (f) PDE residual computation and loss functions

#### (i) Temporal derivatives

Autograd (STAT5): 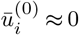. Finite difference (Sturis):

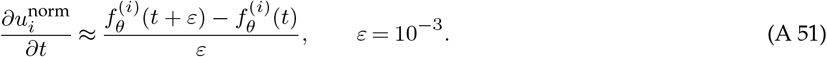

#### (ii) PDE residual

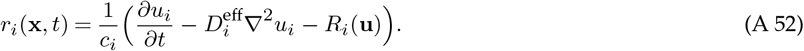

#### (iii) Composite loss function

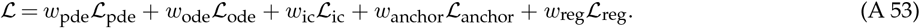

The PDE loss uses Smooth L1 (Huber, *β*_*H*_ = 1.0) with smooth capping:

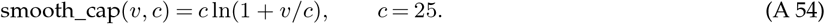

The ODE supervision loss compares spatial means to the ODE reference:

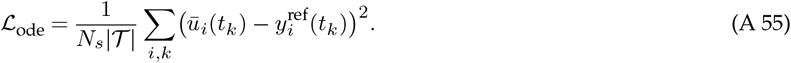

The trajectory anchor loss prevents drift:

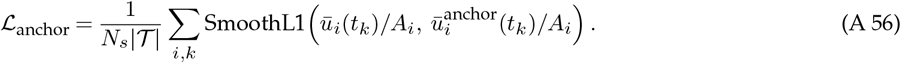

In the L-BFGS and refinement stages, a hybrid loss with species weights (*w*_pApA_ = *w*_pBpB_ = 1.5, *w*_nucpApB_ = 2.5, others = 1.0) and piecewise relative Huber terms is employed.

### (g) Multi-stage training pipelines

#### (i) STAT5 model: three-arm experiment design

##### Arm A: Hidden10 (4-stage pipeline)

*Stage 1: Inverse* (320 steps, Adam, *η* = 2 × 10^−4^): *w*_ode_ = 2.0, *w*_pde_ = 0.15, *w*_ic_ = 1.0. Joint optimization of *θ* and PDE coefficients. *Stage 2: Anchored forward* (320 steps, *η* = 5 × 10^−5^): PDE frozen, *w*_anchor_ = 20.0, *w*_pde_ = 0.02. *Stage 3: L-BFGS* [28] (200 steps, *η* = 1.2 × 10^−4^, history 50, strong Wolfe): *w*_ode_ = 4.0, *w*_anchor_ = 5.0. *Stage 4: AdamW* [29] tail (120 steps, *η* = 10^−5^, weight decay 10^−4^, cosine annealing).

##### Arm B: Strict8 (2-stage pipeline)

Eight observable species only, absolute-softplus decoding, strict relative *ℓ*^2^ loss. Stage 1: Inverse (320 steps). Stage 2: Forward (320 steps, *w*_ode_ = 0.5, PDE frozen).

##### Arm C: PDE-only forward (1 stage)

400 steps, *w*_pde_ = 0.5, *w*_ic_ = 5.0, *w*_ode_ = 0. Frozen PDE coefficients from Arm A.

#### (ii) Ultradian model: three-arm pipeline

##### Arm A: Full PINN (4 stages)

*Stage A*.*1: Inverse* (400 steps, AdamW, *η* = 2 × 10^−4^): PDE warmup over 80 steps, time batch = 2. *Stage A*.*2: Forward* (400 steps): PDE frozen, *w*_anchor_ = 10.0. *Stage A*.*3: L-BFGS* (200 steps, *η* = 5 × 10^−5^). *Stage A*.*4: AdamW tail* (120 steps, *η* = 10^−5^).

##### Arm B: PDE-only forward

*w*_pde_ = 0.30, *w*_ic_ = 5.0, *w*_ode_ = 0, 400 steps.

##### Arm C: ODE with learned rate scales

Direct `scipy.integrate.solve_ivp` (BDF/Radau) using learned rates. No neural network.

#### (iii) Divergence suppression: two-phase PINN training

**Phase 1: Warmstart** (1000 steps, Adam, *η* = 2 × 10^−3^): fit a perturbed ODE with 200 dense points.

**Phase 2: Fine-tuning** (2000 steps, Adam, *η* = 5 × 10^−4^):

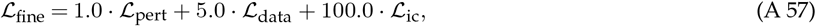

where 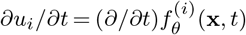.

### (h) Dynamical and parametric sensitivity analysis

Analyses use the 10-species ODE with PINN-learned parameters, [STAT5]_total_ = 1.0, **u**^(0)^ = (0, 0, 1, 0, 0, 1, 0, 0, 1, 0.02)^⊤^.

#### (i) Lyapunov exponent (Benettin method)

Eight tangent vectors (∥***ξ***∥ = 10^−8^), *T* = 4800 min, *N*_renorm_ = 200 segments [33]:

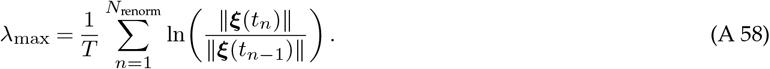

#### (ii) Parameter perturbation study

Five parameters (*k*_phos_, *k*_socs,inh_, *k*_jak,act_, *k*_exp,homo_, *k*_nuc_dephos,homo_), six levels (*δ* ∈ {±0.1%, ±1%, ±5%}). Divergence: *d*(*t*) = ∥**y**_base_(*t*) − **y**_pert_(*t*)∥_2_.

#### (iii) Bifurcation analysis

*k*_socs,inh_ × [0.2, 8.0], 80 values. Record min/max of STAT5A, RJ_active_, SOCS over the last 20% of trajectory.

#### (iv) Data-anchored divergence-suppression protocol

Experimental matrix: 5 parameters × 2 perturbations (+5%, +20%) × 4 data levels (*n* ∈ {8, 16, 32, 64}) = 40 conditions. *T* = 2400 min. Metrics:

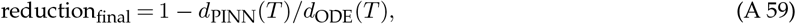

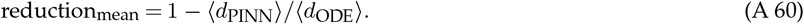

## Software and computational resources

All neural networks are implemented in PyTorch 2.x with CUDA. ODE integrations use `scipy.integrate.solve_ivp` (Radau/BDF). Spectral Laplacians use `torch.fft`. All STAT5 and divergence-suppression experiments fix the global random seed to 42, and computations were performed on an NVIDIA DGX Spark. The complete implementation— entry scripts for the three STAT5 arms (A/B/C), the ultradian insulin–glucose model and the divergence-suppression analyses, together with trained weights and the plotting routines that regenerate every figure and table—is organized as a set of experiment scripts plus result directories and a pinned `requirements.txt`, and is publicly available at https://github.com/xingyang688/PINN-BIO (see Data Accessibility). Each script writes its trajectories, per-species RMSE reports and figures to an `outputs/` directory, so every reported result can be reproduced from the corresponding script under the grids, horizons and step counts specified in Section (g).

## Ethics

This work is entirely computational and did not involve human participants, human data, or animal subjects; no ethical approval was required.

## Data Accessibility

All code, trained model weights and the scripts that reproduce every figure and table in this study are openly available under the MIT license in a public GitHub repository (https://github.com/xingyang688/PINN-BIO) and are archived— together with the trained weights and a README documenting the exact reproduction commands—in a permanent Zenodo record (https://doi.org/10.5281/zenodo.XXXXXXX) [34]. No experimental data were generated; all reference trajectories are produced by the deposited code. The Zenodo DOI will be finalized at the revision stage.

## Authors’ Contributions

Contributions follow the CRediT taxonomy. **J.D**.: conceptualization, formal analysis, investigation, writing— review and editing. **X.Y**.: methodology, software, validation, visualization, supervision, project administration, funding acquisition, writing—review and editing. **Xuchang Z**. and **Xinyu Z**.: writing—original draft, writing—review and editing. All authors gave final approval for publication and agree to be held accountable for the work performed therein.

## Competing Interests

The authors declare that they have no competing interests.

## Funding

This work was supported by the National Natural Science Foundation of China (grant No. 12464009) and the Natural Science Foundation of Guangxi (grant No. AD21220127).

## Disclaimer

During the preparation of this work, the authors used Grok 4.1 by xAI to assist with language editing and manuscript refinement. After using this tool, the authors reviewed and edited the content as needed and took full responsibility for the publication’s content.

## Notes

### Competing Interest Statement

The authors have declared no competing interest.

## References

1 A. M. Turing, “The chemical basis of morphogenesis”, Philos Trans R Soc Lond B Biol Sci 237, 37–72 (1952).

2 A. Goldbeter, Biochemical Oscillations and Cellular Rhythms (Cambridge University Press, 1996).

3 M. Raissi, P. Perdikaris, and G. E. Karniadakis, “Physics-informed neural networks: A deep learning framework for solving forward and inverse problems involving nonlinear partial differential equations”, J Comput Phys 378, 686–707 (2019).

4 J. H. Lagergren, J. T. Nardini, R. E. Baker, M. J. Simpson, and K. B. Flores, “Biologically-informed neural networks guide mechanistic modeling from sparse experimental data”, PLoS Comput Biol 16, e1008462 (2020).

5 C. Rao, H. Sun, and Y. Liu, Physics-encoded recurrent convolutional neural network for solving spatiotemporal PDEs, (2021)

6 Z. Chen, Y. Liu, and H. Sun, “Physics-informed learning of governing equations from scarce data”, Nat Commun 12, 6136 (2021).

7 B. Bezekçi, “The refined physics-informed neural networks for nonlinear convection–reaction–diffusion equations using exponential schemes”, Black Sea J Eng Sci 8, 10.34248/bsengineering.1645207 (2025).

8 G. E. Karniadakis, I. G. Kevrekidis, L. Lu, P. Perdikaris, S. Wang, and L. Yang, “Physics-informed machine learning”, Nat Rev Phys 3, 422–440 (2021).

9 S. Vandvajdi, Y. Mao, M. Poudineh, and M. Kohandel, “Investigating glucose–lactate metabolism in glioblastoma multiforme via universal physics-informed neural networks”, Math Biosci Eng 22, 2486–2505 (2025).

10 D. Lao-Martil, J. P. J. Schmitz, B. Teusink, and N. A. W. van Riel, “Elucidating yeast glycolytic dynamics at steady state growth and glucose pulses through kinetic metabolic modeling”, Metab Eng 77, 128–142 (2023).

11 J. Ul Rahman, S. Danish, and D. Lu, “Deep neural network-based simulation of the Sel’kov model in glycolysis: a comprehensive analysis”, Mathematics 11, 3216 (2023).

12 E. E. Sel’kov, “Self-oscillations in glycolysis. 1. A simple kinetic model”, Eur J Biochem 4, 79–86 (1968).

13 O. Lo-Thong, P. Charton, X. F. Cadet, B. Grondin-Perez, E. Saavedra, C. Damour, and F. Cadet, “Identification of flux checkpoints in a metabolic pathway through white-box, grey-box and black-box modeling approaches”, Sci Rep 10, 13446 (2020).

14 A. Akbari, Z. B. Haiman, and B. O. Palsson, “A data-driven approach for timescale decomposition of biochemical reaction networks”, mSystems 9, e01001–23 (2024).

15 B. Huard, A. Bridgewater, and M. Angelova, “Mathematical investigation of diabetically impaired ultradian oscillations in the glucose–insulin regulation”, J Theor Biol 418, 66–76 (2017).

16 J. Li, Y. Kuang, and C. C. Mason, “Modeling the glucose–insulin regulatory system and ultradian insulin secretory oscillations with two explicit time delays”, J Theor Biol 242, 722–735 (2006).

17 A. Bridgewater, B. Stringer, B. Huard, and M. Angelova, “Ultradian rhythms in glucose regulation: a mathematical assessment”, AIP Conf Proc 2090, 050010 (2019).

18 S. Ruschel and B. Huard, Sounding the metabolic orchestra: a delay dynamical systems perspective on the glucose–insulin regulatory response to on–off glucose infusion, (2023)

19 M. Daneker, Z. Zhang, G. E. Karniadakis, and L. Lu, Systems biology: identifiability analysis and parameter identification via systems-biology informed neural networks, (2022)

20 S. De Carli, N. Licini, D. Previtali, F. Previdi, and A. Ferramosca, Integrating biological-informed recurrent neural networks for glucose–insulin dynamics modeling, (2025)

21 V. Roquemen-Echeverri, T. Kushner, P. G. Jacobs, and C. Mosquera-Lopez, A physiologically-constrained neural network digital twin framework for replicating glucose dynamics in type 1 diabetes, (2025)

22 A. Yazdani, L. Lu, M. Raissi, and G. E. Karniadakis, “Systems biology informed deep learning for inferring parameters and hidden dynamics”, PLoS Comput Biol 16, e1007575 (2020).

23 M. de Rooij, B. Erdoős, N. A. W. van Riel, and S. D. O’Donovan, “Physiology-informed regularisation enables training of universal differential equation systems for biological applications”, PLoS Comput Biol 21, e1012198 (2025).

24 C. Rackauckas, Y. Ma, J. Martensen, et al., Universal differential equations for scientific machine learning, (2020)

25 J. Sturis, K. S. Polonsky, E. Mosekilde, and E. Van Cauter, “Computer model for mechanisms underlying ultradian oscillations of insulin and glucose”, Am J Physiol Endocrinol Metab 260, E801–E809 (1991).

26 M. E. Boehm, L. Adlung, M. Schilling, S. Roth, U. Klingmüller, and W. D. Lehmann, “Identification of isoform-specific dynamics in phosphorylation-dependent STAT5 dimerization by quantitative mass spectrometry and mathematical modeling”, J Proteome Res 13, 5685–5694 (2014).

27 S. Wang, X. Yu, and P. Perdikaris, “When and why PINNs fail to train: A neural tangent kernel perspective”, J Comput Phys 449, 110768 (2022).

28 D. C. Liu and J. Nocedal, “On the limited memory BFGS method for large scale optimization”, Math Program 45, 503–528 (1989).

29 I. Loshchilov and F. Hutter, “Decoupled weight decay regularization”, in International Conference on Learning Representations (ICLR) (2019).

30 K. Linka, A. Schäfer, X. Meng, Z. Zou, G. E. Karniadakis, and E. Kuhl, “Bayesian physics-informed neural networks for real-world problems”, Comput Methods Appl Mech Eng 402, 115346 (2022).

31 D. Hendrycks and K. Gimpel, Gaussian Error Linear Units (GELUs), (2016)

32 J. L. Ba, J. R. Kiros, and G. E. Hinton, Layer normalization, (2016)

33 G. Benettin, L. Galgani, A. Giorgilli, and J. M. Strelcyn, “Lyapunov characteristic exponents for smooth dynamical systems and for Hamiltonian systems; a method for computing all of them. Part 1: Theory”, Meccanica 15, 9–20 (1980).

34 X. Yang, J. Deng, X. Zhang, and X. Zhang, PINN-BIO: Physics-informed neural networks for 3D diffusion–reaction modeling of biochemical dynamics (code and trained weights), https://github.com/xingyang688/PINN-BIO, version v1.0, Placeholder DOI; to be minted from the GitHub release at revision, 2026.

35 L. Lu, X. Meng, Z. Mao, and G. E. Karniadakis, “DeepXDE: A deep learning library for solving differential equations”, SIAM Rev 63, 208–228 (2021).

36 W. Ji, W. Qiu, Z. Shi, S. Pan, and S. Deng, “Stiff-PINN: Physics-informed neural network for stiff chemical kinetics”, J Phys Chem A 125, 8098–8106 (2021).

